# Healthy ageing delays the neural processing of face features relevant for behaviour by 40 ms

**DOI:** 10.1101/326009

**Authors:** Katarzyna Jaworska, Fei Yi, Robin A.A. Ince, Nicola J. van Rijsbergen, Philippe G. Schyns, Guillaume A. Rousselet

**Affiliations:** Institute of Neuroscience and Psychology, University of Glasgow, Glasgow, United Kingdom

## Abstract

Fast and accurate face processing is critical for everyday social interactions, but it declines and becomes delayed with age, as measured by both neural and behavioural responses. Here, we addressed the critical challenge of understanding how ageing changes neural information processing mechanisms to delay behaviour. Young (20-36 years) and older (60-86 years) adults performed the basic social interaction task detecting a face vs. noise while we recorded their electroencephalogram (EEG). In each participant, using a new information theoretic framework we reconstructed the features supporting face detection behaviour, and also where, when and how EEG activity represents them. We found that occipital-temporal pathway activity dynamically represents the eyes of the face images for behaviour ∼170 ms post-stimulus, with a 40 ms delay in older adults that underlies their 200 ms behavioural deficit of slower reaction times. Our results therefore demonstrate how ageing can change neural information processing mechanisms that underlie behavioural slow down.

**Author summary:** Older adults are consistently slower than young adults in a variety of behavioural perceptual tasks. So far, it has been unclear if the underlying cause of the behavioural delay relates to attentional or perceptual differences in encoding visual information, or slower neural processing speed, or other neural factors. Our study addresses these questions by showing that in a basic social interaction task (discriminating faces from noise), young and older adults encoded the same visual information (eyes of the face) to perform the task. Moreover, early brain activity (within 200 ms following stimulus onset) encoded the same visual information (again, eyes of the face) in both groups, but was delayed and weaker in older adults. These early delays in information encoding were directly related to the observed behavioural slowing in older adults, showing that differences in early perceptual brain processes can contribute to the motor response.

## Introduction

There is strong evidence that an age-related slowing down occurs when performing various behavioural tasks (1). There is also parallel evidence that age-related changes in the brain can slow down neural processing (2–6). Although these studies can inform where and when, in the brain, ageing can impact neural activity, a critical challenge remains to develop theories of cognitive ageing that explain how the neural information processes that are involved in a cognitive task contribute to slow down behaviour. It is imperative to address this challenge to understand how ageing impacts the specific cognitive mechanisms that mediate behaviour. To illustrate, consider the fundamental social cognition task of detecting a face (see Figure 1). Although we expect older participants to detect faces in this task more slowly, a more complete understanding of the effects of aging on cognitive processing requires richer data. At a minimum, we need to characterize the face information that young and older participants selectively use when detecting faces, for example, because slower face detection could result from older adults needing more facial features, or simply because older participants do not use the same features. We also need to characterize the information content of the neural activity underlying the behavioural task at hand, because the age-related slowing could arise from neural representation of several task-irrelevant features (a decline of selectivity). Finally, we would also need to trace the origin of the behavioural delay in the neural mechanisms that represent the face for the detection task *per se*, for example, because older brains could be generally delayed in their onset of neural activity when responding to any visual stimuli, rather than specifically delayed when processing the task-relevant face features.

**Figure 1.**
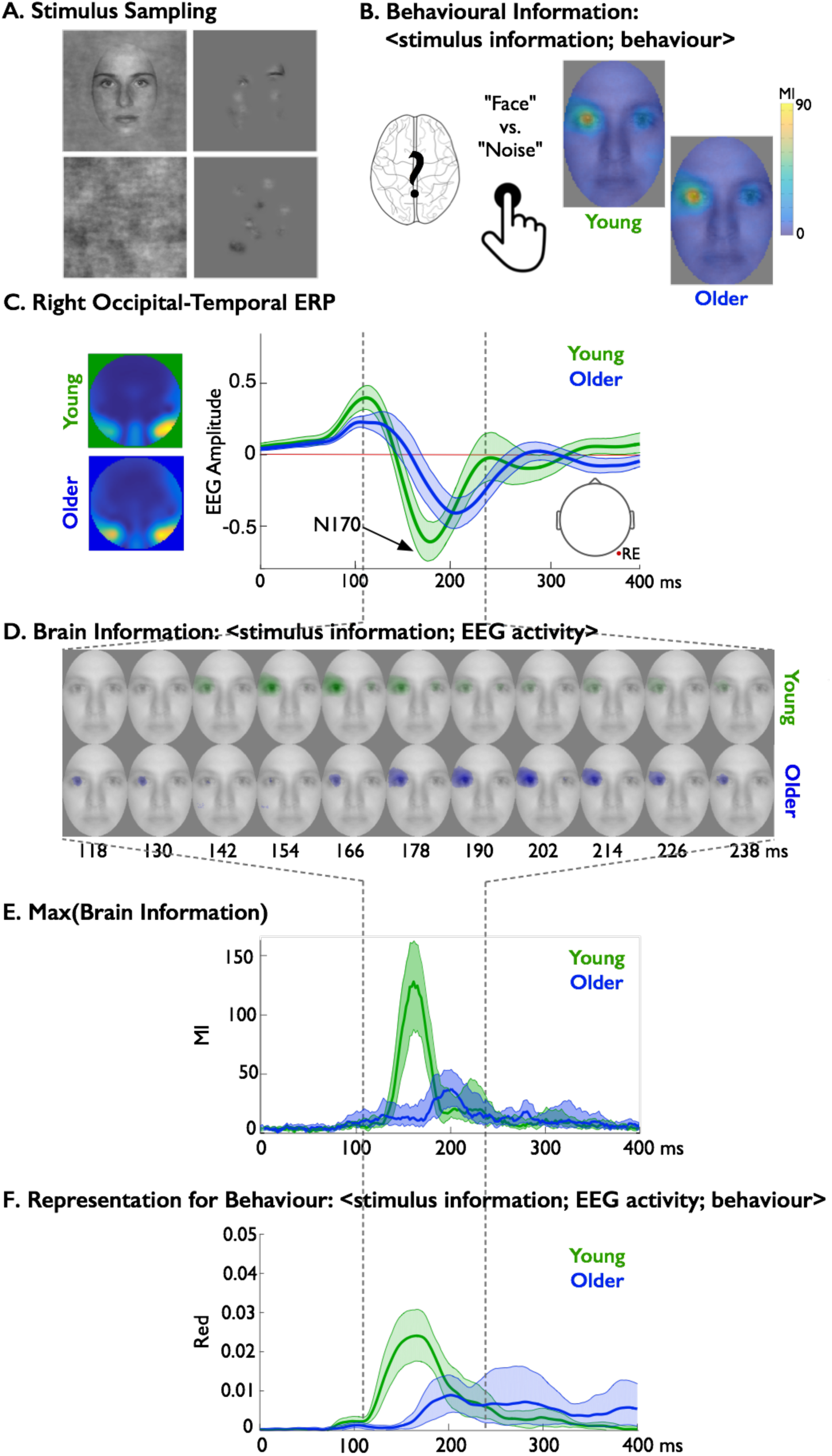
Illustration of the experimental paradigm and analyses. **(A) Stimuli.** Eighteen young and nineteen older participants each detected faces vs. noise textures from 2,220 pictures that revealed information through Gaussian apertures (“Bubbles” (7)), while we recorded their face detection behaviour (with key presses) and EEG brain activity. We performed the following computations in each individual observer. **(B) Behavioural information**. We used MI to compute <stimulus information; behaviour>, the relationship between stimulus pixel visibility on each trial and the corresponding reaction times. The same left eye of faces was associated with faster reaction times in both young and older participants. **(C) Event Related Potentials (ERPs).** Based on these trials, a full brain analysis revealed larger average ERPs associated with face detection at the right hemisphere occipital-temporal electrode (RE, whose location is sketched with a cranial view of the head underneath) of both young and older adults. **(D) Brain information**. Using MI, we computed <stimulus information; EEG activity> every 2 ms post stimulus, which represents the relationship between stimulus pixel visibility in a single trial and the corresponding electrode voltage amplitudes, to reveal the dynamics of any visual feature represented in the variations of the EEG. For illustration, we plotted examples of these classification images every 12 ms between 118 and 238 ms post stimulus. The resulting MI images revealed that the N170 of both young and older adults represented the same left eye, although older adults did so with a delay. **(E) Max(Brain Information).** To precisely estimate this age-related, feature-processing delay, we plotted the time course of maximum MI (across the pixels of each MI image in **(D)**) between 0 and 400 ms post stimulus, which peaked 40 ms later in older participants. **(F) Representation for behaviour.** Finally, we confirmed that these features are represented in the brain to support face detection behaviour. To do this, we computed feature redundancy (*FeatRed*), which quantifies the common effect of stimulus information on both EEG activity and behaviour from the triple relationship <stimulus information; EEG activity; behaviour>.

In this study, we present a new paradigm to advance and deepen our understanding of information processing in cognitive aging. We developed Stimulus Information Representation (SIR), an information theoretic framework, to tease apart stimulus information that supports behaviour from that which does not. This new framework considers the interactions between three important variables, not just two, as is the norm: stimulus information, neural EEG activity and behaviour. We used the Bubbles technique (7) to randomly sample visual information from the stimulus on each trial (see Figure 1A), which limits researcher bias by not making a priori assumptions about the features that participants in different groups use for the task. With the double relationship <stimulus information; behaviour>, we coupled the stimulus information sampled on each trial with the corresponding participant detection behaviour to reconstruct the features that underlie their detection behaviour (see Figure 1B). With the double relationship <stimulus information; EEG activity>, we coupled the same sampled information with the corresponding EEG activity of each participant’s brain engaging with the detection task, to independently characterize where, when and how brain activity represents face features (see Figure 1C-E). Finally, with the new triple relationship <stimulus information; EEG activity; behaviour>, we directly visualized with a single integrated measure the dynamic development of the representation of a face in the brain to support face detection behaviour (see Figure 1F). We computed the dynamics of representation individually in 17 young and 18 older participants and showed, using information theoretic redundancy across participants, that delayed behavioural information processing and delayed reaction time reflect a common ageing factor within our sample. We now present each double relationship in turn, followed by the triple relationship, and the group redundancy analysis. Section titles indicate the specific relationship that we evaluate between the three variables.

## Results

### A. Behavioural information: <stimulus information; behaviour>

Seventeen young (median age = 23) and eighteen older (median age = 66) participants categorized 2,200 pictures of faces and noise textures revealed through Gaussian apertures (so-called “Bubbles” (7)), which sample random spatial regions of face and noise images on each trial. Young participants were 198 ms faster to detect faces (median reaction time (RT) in young = 378 ms, 95% confidence interval = [349, 401] vs. RT in older participants = 576 ms [527, 604]). To determine the relationship between sampled face information (pixels) and the varying RTs on each trial, we computed MutuaI Information (MI) between <stimulus information; RT> (see *Materials and Methods*). We found that the presence of the left eye in image samples modulated RTs in all young participants (17/17) and in most older participants (16/18; see Figure 2). The presence of the right eye also modulated RTs in a few young and older participants. However, whereas young participants could use many other features to perform the task accurately, older participants specifically used the eyes (see *Supplementary Results* and *Supplementary Figures S1-S4*).

**Figure 2.**
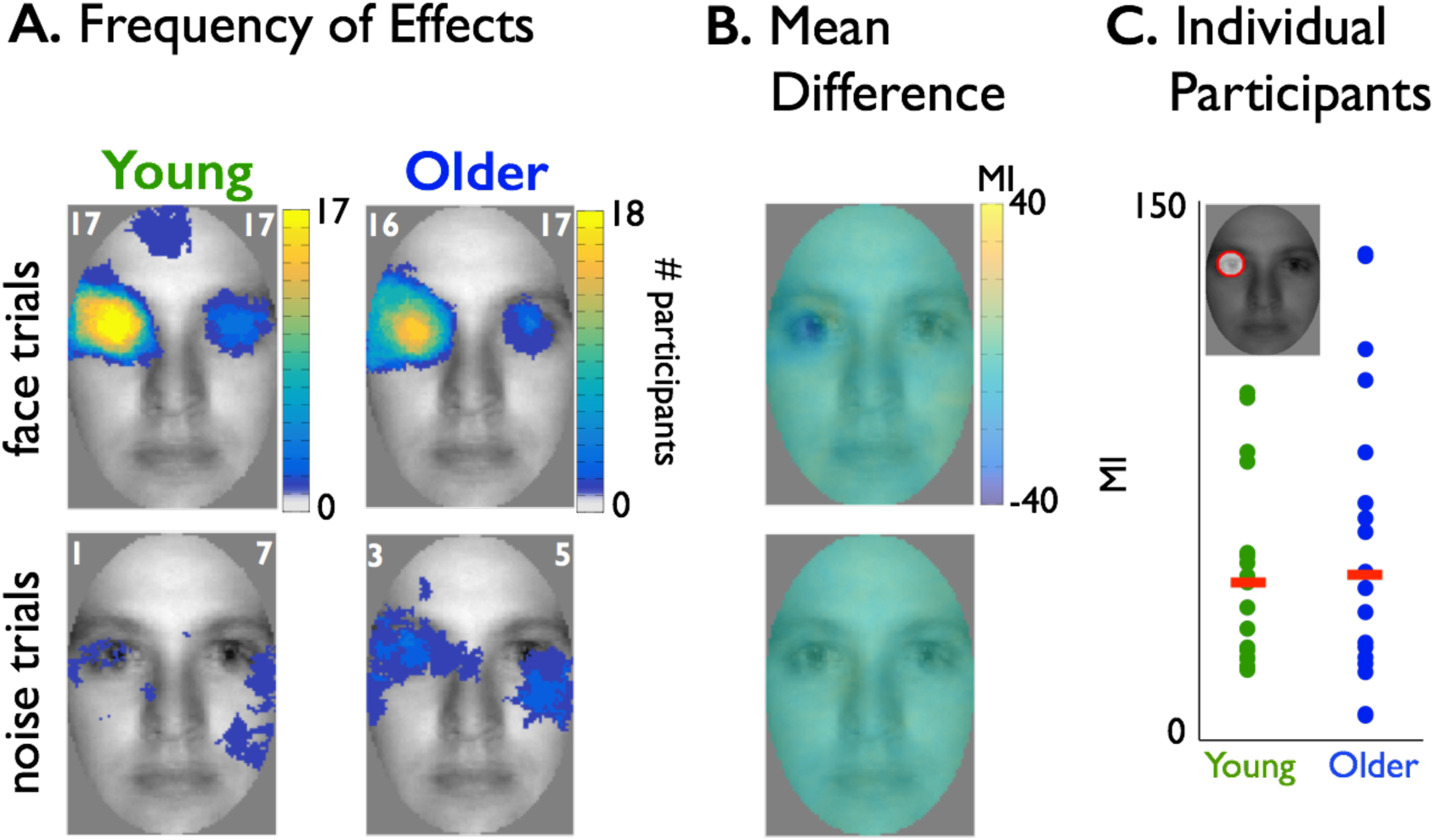
Behavioural Information. (A) Significant effects. The white number in the left (vs. right) upper corner of each face image indicates the maximum number of participants showing a significant relationship between pixel visibility and RTat the same (vs. any) face pixel. Yellow colours reflect that a high number of significant effects across participants cluster in the left eye region of the image in young and older participants. As such, for most young and older participants, the left eye of the face image was associated with faster RTs (see also *Supplementary Results*). **(B) Mean difference**. Images show the differences between average MI images (young – older) for face trials (top) and noise trials (bottom). The scale represents normalised MI values (see *Materials and Methods*). **(C) Individual Participants**. Dot plot shows, for each participant (coloured dot), MI values summed within a mask that captures left eye pixels (represented as a red circle in the face inset; see *Supplementary Methods*). Red bars correspond to the median of these per-participant averages. Distributions of individual participant values were similar between the two groups.

### B. EEG face information representation: <stimulus information; EEG activity>

We now turn to the important question of where, when and how brain processes represent task-relevant features to support behavioural decisions. Before we proceed, we must first rule out low-level optical factors as the main contributor to any age-related delay in our analyses. To do this, we computed the time course of the standard deviation of the mean ERP across electrodes (ERP_STD_) using causal-filtered data (see *Materials and Methods* and Figure 3A). ERP_STD_ onsets correspond to the initial activation of the occipital cortex that follows stimulus presentation (8), which could reveal generic, age-related delays already present in the earliest stages of the visual processing pathway. However, we found similar ERP_STD_ onsets in young (68 ms [64, 72]) and older (69 ms [62, 75]) participants (Figure 3A), with a negligible difference between them (−0.5 ms [-7, 5]), which suggests that there is no clear evidence for a generic delay in the onset of cortical activity in response to visual stimuli in older participants.

**Figure 3.**
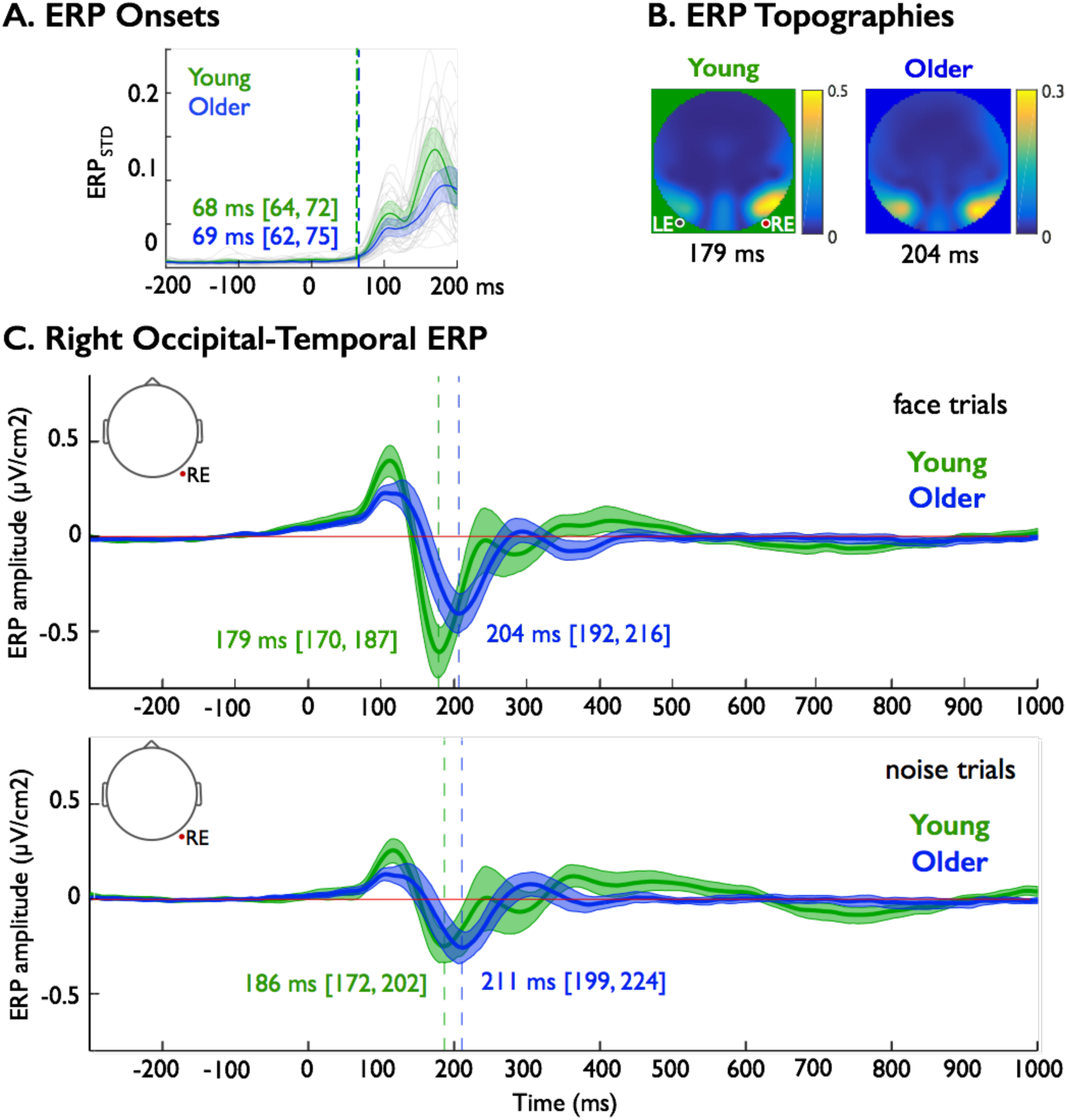
N170 Event Related Potentials. (A) ERP Onsets. Thin grey lines show individual participants’ ERP_STD_ (µV/cm^2^). Coloured thick lines show group averages with shaded areas indicating 95% confidence intervals around the group means. Vertical dashed lines mark the overlapped onset times of cortical activity in each group. **(B) ERP Topographies.** For face only trials, we averaged across participants the largest squared ERP amplitude at the time of its peak. **(C). Right Occipital-Temporal ERP.** Thick lines show averaged ERPs across young (green) and older (blue) participants, for face trials (top panel) and noise bubble trials (bottom panel), with shaded areas corresponding to 95% confidence intervals. In each panel, numbers indicate the group median (and confidence intervals) of N170 latencies, which are delayed by ∼20 ms in older adults.

#### B.1. N170 is delayed in older participants

Despite no clear generic neural delay in older participants, their peak N170 response latencies to full faces on the right occipital electrode (RE) were delayed relative to the N170 response of young participants by 18 ms [9, 24] (and by 23 ms [9, 38], in response to the practice full noise trials; see Figure 3B for a summary topography, see Materials and Methods, Stimuli). These N170 peak delays on the RE were confirmed in bubbles trials that sampled face and noise information (i.e. 22 ms [10, 32] and 18 ms [7, 31], respectively, see Figure 3C). N170 peak amplitudes were similar in young and older participants, except on practice noise trials, where they were larger in older than young participants (see *Supplementary Results* and *Supplementary Figure S5*).

#### B.2. Contra-lateral eye representation over the N170 time course

Our novel methodology enables us to directly visualize the dynamic representation of facial features in the single-trial amplitude variations of EEG activity during the detection task. To do this, we computed the MI relationship between <pixel visibility information; EEG activity>, for each stimulus pixel, on the left (LE) and right (RE) electrode (see *Materials and Methods, Electrode Selection*), every 2 ms between 0 and 400 ms post-stimulus. Figure 1D illustrates example MI classification images of brain information on electrode RE.

As shown in Figure 4A, EEG activity at RE contra-laterally represented the left eye in most young (N = 16/17) and older (N = 12/18) adults (see *Supplementary Figure*s *S7-S8* for LE results), with a weaker overall representation in older adults (their peak MI was 57% [42, 82] of that of young adults, see Figure 4B-C).

**Figure 4.**
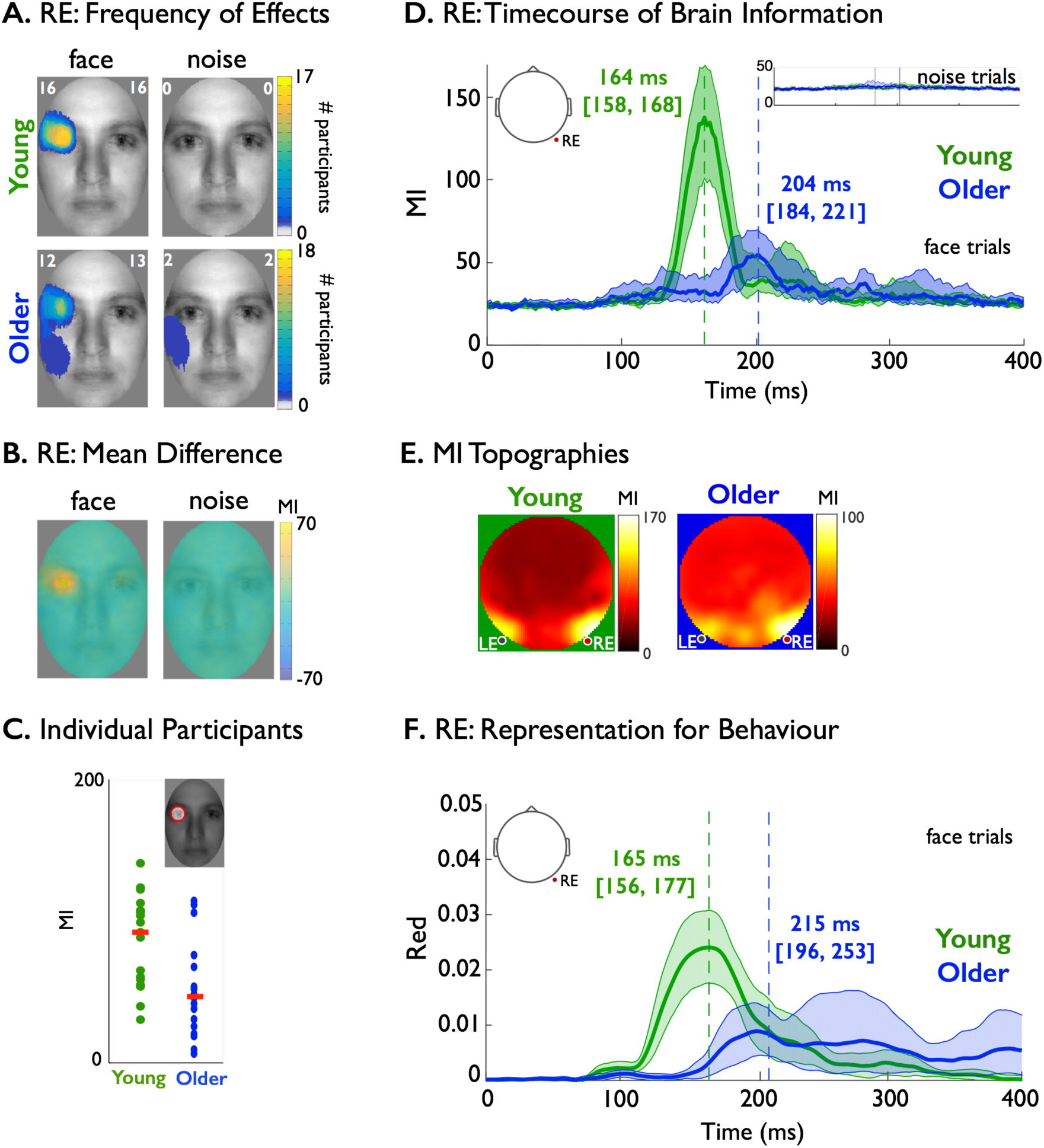
Brain Information. (A) RE: Frequency of Significant Effects. Using the maximum MI image across time points on RE in each participant, the white number in the left upper corner of every image corresponds to the maximum number of participants showing a significant effect at the same face pixel (on face and noise trials separately), whereas the number in the right upper corner corresponds to the total number of participants showing significant effects at any pixel. **(B) RE: Mean Difference**. Averaging the maximum MI images across young and older participants separately produced images of group differences (young – older) for face trials (left) and noise trials (right). Average MI in the left eye region of the face image was higher for young than older participants. Throughout the figure, MI represents normalised MI values (see *Materials and Methods*). **(C) Individual Participants.** Dot plot shows, for each participant (coloured dot), MI values summed within a mask that captures left eye-only pixels (represented as a red circle in the face inset; see *Supplementary Methods*). Red bars correspond to the median of these per-participant averages. Distributions of individual participant values were different between the two groups. **(D) RE: Timecourse of Brain Information** are presented for both face and noise (insets) bubbles trials on RE. Colour-coded numbers correspond to median latencies of maximum MI in both groups. Shaded areas correspond to bootstrap 95% confidence intervals around the 20% trimmed mean. **(E) MI Topographies.** MI<face information; EEG activity> (i.e. brain information) averaged across young and older participants for each electrode across time points and face pixels**. (F) RE: Representation for Behaviour.** A time course of behavioural redundancy shows a 46 ms delay in the representation of the left eye on the RE for delayed face detection reaction time in older adults.

We then used the resulting images of brain information to compute their maximum MI across all time points and pixel locations (see Figure 4D). Peak EEG representation of the left eye occurred 40 ms [23, 57] earlier in young adults relative to older adults (i.e. ∼164 ms in young vs. ∼204 ms in older adults).

To rule out the possibility that maximum information was represented on other electrodes, we independently performed a full brain-by-time analysis. To this end, we again computed the MI relationship between <pixel visibility information; EEG activity> at each pixel and time point between 0 and 400 ms post-stimulus, on all electrodes. We found that maximum MI peaked primarily in the left and right lateral-occipital region in both young and older observers (see Figure 4E). Furthermore, computing the maximum MI across all electrodes revealed similar results to those reported here for RE (see *Supplementary Figure S6*).

At this juncture, it is important to emphasize that we also ruled out a potential effect of participants paying spatial attention to the location of the eyes in sample images (rather than one of representation of the eyes per se) by repeating the above analyses using only noise trials. In these noise trials, we did not find sensitivity to the image locations of the eyes (i.e. where the eyes of a face would be in face trials), in any participant. Note that the midline electrode (Oz, see *Supplementary Figures S7-S8*) also revealed the weaker representation of various other facial features (such as eyes, chin, mouth, nose, and forehead) in some participants. As these representations were inconsistent across participants, we did not analyse them further.

### C. Stimulus representations in the brain to support behavioural decision: <stimulus information; EEG activity; behaviour>

So far, we have shown that face detection behaviour in all participants involves the processing of eyes in the stimulus (particularly the left one), with a ∼200 ms decision response delay in the older group. We have also shown that this left eye impacted neural EEG responses in most participants, with a 40 ms representation delay in the older group. Now, we integrate behavioural and brain results by showing where the representation of the eyes in the EEG also modulates face detection behaviour, on the same trials.

To do this, we computed separately for each participant the information theoretic feature redundancy (9), Red(eye; EEG; RT), which is the shared variability between three variable: single-trial visibility of the left and right eye in the stimulus, at contralateral electrodes (RE and LE), together with the corresponding RTs (see *Materials and Methods, Feature Redundancy*). In this application, redundancy measures the strength of the representational similarity of a given stimulus feature (e.g. relative visibility of the left eye) between variations of EEG amplitude (e.g. on the RE) and variations of face detection RT. On the RE, we found a 46 ms [30, 81] redundancy delay between the young and older participants (i.e. 165 ms peak [155, 178] for young, and a 215 ms peak [196, 246] for older adults; see Figure 4F). We observed a similar 42 ms [20, 86] delay between young and older participants on the LE, where right eye redundancy peaked at 172 ms [161, 186] for young, and at 220 ms [196, 261] for older adults (see *Supplementary Figure S10*). These results confirm that the EEG and RT similarly represent the eyes, but with a 42 ms delay in older adults.

### D. Delayed stimulus representations in the brain underlie delayed behavioral reaction times

We now demonstrate that the delayed representation of the eyes in the EEG underpins delayed RT in older adults. Using SIR, we tested this directly with redundancy at group level, by computing in two steps the relationship between the three variables (young vs. older adults, peak feature redundancy time, and median RT; see *Materials and Methods, Group Redundancy*). In the first step, we quantified (with MI) the strength of the relationship between age group and each participant’s median RT – it was 0.5 [0.36 0.81] bits – and also between age group and redundant bilateral eye representation latency in the EEG – it was 0.48 [0.34 0.74] bits. Having shown that both RT latency and EEG eye representation correlate with age group with similar strengths, in the second step we asked whether their correlations overlap – as would happen if a delayed EEG representation was systematically related to a delayed RT delay across participants of both age groups. Indeed, we found that the latency of left eye representation at RE is 100% redundant with RT – 0.28 [0.06 0.51] bits – whereas the right eye (at LE) is only 40% redundant with RT – 0.12 [-0.09 0.43] bits. This demonstrated that only the left eye (at RE) representation is purely related to the RT deficit between age groups. These results reflect the strongly lateralized relationship between stimulus and RT (Figure 2).

To summarize, we used information theoretic redundancy at two levels (within and across participants) to demonstrate a direct link between behaviourally relevant neural information processing change (48 ms delay in left eye redundancy) and a behavioural ageing deficit (200 ms reaction time delay across participants of our sample). Specifically, within participants, we explicitly quantified the triple relationship <stimulus information; EEG activity; behaviour> within SIR to focus on the aspects of information processing reflected in EEG signals that are directly relevant to face detection RT. Across participants, quantification of the triple relationship <age group; representational delay; RT delay> suggests a group level interpretation in which neural representational delays underlie behavioral slow down in ageing.

## Discussion

Here, we set out to investigate how information-processing delays in the ageing brain could slow down face detection behaviour (a fundamental social interaction task), using the novel information theoretic measures of the SIR framework. Specifically, we considered the interactions between three variables: stimulus information, neural EEG activity, and reaction time, and compared the results between young and older adults. We characterized the face information that young and older participants selectively use when detecting faces (i.e. the eyes of a face), and we traced the origin of the behavioural delay with ageing to a delay in the neural processes that represent the eyes of faces for the task during the N170 period. We ensured that the neural eye representation delay could not be explained by generic delays of onsets of cortical activity in young and older participants.

This study provides an important step towards understanding visual cognitive ageing. First, we demonstrate that young and older adults can use the same face information (the eyes) to quickly detect faces. As such, our results contrast with previously published findings that suggest that a differential use of horizontal vs. vertical information might underlie the impairment of older adults when identifying faces (10, 11). Here, although older adults had fewer correct responses when detecting faces, the same face information modulated their slower reaction times. Our analyses of correct vs. incorrect responses (see *Supplementary Results*) revealed that while young adults can use any facial features to detect a face, older adults relied heavily on the eyes for correct detection. As such, it is possible that rather than inefficiently extracting information from faces, older adults were more conservative, detecting a face only when its eyes were visible (see also (12)), although another possibility is that older adults relied more on local contrast information contained within the eye region of the face, in line with previous studies showing that they require more contrast to detect and discriminate faces (13, 14). In any case, the uncovering of such difference was made possible by Bubbles sampling (7), which limits researcher bias by not making a priori assumptions about the features that participants should use (12,15,16).

Establishing equivalence of behavioural information in the two groups is an important benchmark for comparing the task-relevant neural coding of information. Here, we found that the EEG activity of both young and older adults represented the eye pixels contralateral to the lateral-occipital recording electrodes, an effect that was stronger at the right hemisphere electrodes in both groups, in agreement with the reported right hemisphere dominance for face processing (17). Although feature representation was qualitatively similar across young and older adults, it was delayed by 40 ms and weaker in older adults, in the absence of generic delays in the onset of visual cortical activity in older participants. This suggests that the reported delay occurred at the stages of cortical information processing, and was not due to pre-cortical neural factors; thereby adding to the evidence that processing speed delays are unlikely to be due to bottom-up optical factors, such as senile miosis, contrast sensitivity (18, 19), or visual acuity (20). Furthermore, we believe that bottom-up optical factors were unlikely contributors to the observed differences at the neural level, because any bottom-up factors should affect all neural responses irrespectively of their category, whereas we observed much larger N170 to noise textures in older than in young participants.

The age-related delay that we report here agrees with previous cross-sectional results, which suggest that face processing slows down from 20 years of age and onwards (2,3,18). However, previous studies could not ascribe these delays to the representation of task-relevant features, as reported here (see also (21–23)). Such task-relevant feature representations in the brain are important for two main reasons. First, by demonstrating the neural representation of task-relevant face features in older adults, we can show that the reported delays (neural and behavioural) do not arise from an inability to inhibit the task-irrelevant information that increases with age (4, 24). Second, we can use the direct evidence of neural representation of the same features to demonstrate that the N170 time window is functionally equivalent in young and older adults (cf. (3)). The N170 is an early component of face processing (23,25,26) that is delayed in older participants (2,4–6). Comparing the latencies from the same component across age groups presumes that it indexes the same neuronal processes over a life span. However, prior to this current study, it was unclear whether the single-trial activity evoked during the N170 time window represents the same information content across age groups. Here, we show that the N170 performs the same representation function in two age groups detecting faces, albeit with a delay in older adults (2,21,22,27). In fact, our results suggest that the 40 ms representation delay of the eyes over the N170 in older adults was directly related to their 200 ms reaction time delay when using this information to detect faces. We can thus trace the origin of the behavioural slowing to the early stages of stimulus processing, which could complement other studies that have indicated that slowing originates from motor response generation (28, 29).

At this juncture, it is important to emphasize that the age-related delay in the processing of the eye information from the sample image cannot be attributed to the presence of Bubble masks. Bubbles can be thought of as a ‘masking procedure’ that degrades the visual input and possibly entails object completion (30). The processing of occluded stimuli by the visual system might require additional resources to perform the task, leading to longer processing times (31). As such, any delay observed in a sample of older adults could be due to a combination of factors: a genuine slowing down of processing speed, as well as an increase in the time needed to process the occluded stimulus with respect to young adults. However, our ERP results show that the extra processing time of Bubbled images compared with full images differed very little between young and older participants. Specifically, even though the processing of the Bubbled stimuli was delayed with respect to full images by about 20 ms in both young and older participants, there was only a weak interaction between age and masking condition. In both practice (unmasked) and Bubble (masked) trials, the N170 latency to face images in older participants was delayed by about 20 ms (18 ms in practice trials and 22 ms in Bubble trials) with respect to that in young participants. This agrees with a recent study (18), which showed that even though stimulus luminance affects the entire ERP time course in both young and older participants, it does not affect age-related differences in processing speed.

Future studies should aim to uncover the extent to which delays in other processing stages (i.e. post-perceptual decision, sensorimotor integration, or motor generation processes) influence the observed behavioural slow down. Our results also raise further considerations regarding their generalization to other stimulus categories, both simple and complex, such as scenes, objects, words, but also sensory modalities and tasks. Recent evidence suggests that ageing cannot be regarded as a unitary concept that affects functionally relevant brain regions in the same manner (20). Instead, whether the observed age-related delay is constant or cumulative, may depend on a variety of factors other than structural differences in brain regions (20,32–34), such as the sensory modality, task, and stimulus complexity. Our new methods can address some of these issues by linking tightly controlled stimulus information to brain response and decision behaviour in important social cognition tasks. These methods can also be extended to other sensory modalities to study group differences (i.e. cultural, developmental, clinical) in perception and cognition.

To summarize, our results provide the first functional account that advancing age involves differences in neural delays in the representation of task-relevant information for behaviour. They address the fundamental challenge of developing theories of cognitive ageing that explain how the neural information processes involved in a cognitive task slow down behaviour.

## Materials and Methods

### Participants

Eighteen young (9 females, median age = 23, min 20, max 36) and nineteen older adults (7 females, median age = 66, min 60, max 86) participated in the study (data of fifteen of the young participants were used in (23)). All older adults were local residents. We excluded participants if they reported any current eye condition (i.e., lazy eye, glaucoma, macular degeneration, cataract), had a history of mental illness, were currently taking psychotropic medications or used to take them, suffered from any neurological condition, had diabetes, or had suffered a stroke or a serious head injury. We also excluded participants if their latest eye test was more than a year (for older participants) or two years (for young participants) prior to the study taking place. Two older participants reported having cataracts removed, and one older participant reported having undergone a laser surgery. We included them because their corrected vision was within normal range. We assessed participants’ visual acuity and contrast sensitivity in the lab using a Colenbrander mixed contrast card set and a Pelli-Robson chart. All participants had normal or near-normal visual acuity as measured with the 63 cm viewing distance (computer distance) chart (Table 1). Three older participants had contrast sensitivity of 1.65, and all others had contrast sensitivity of 1.95 log units, within the normal range of contrast sensitivity for that age group (35). All young participants had contrast sensitivity of 1.95 log units or above. During the experimental session, participants wore their habitual correction if needed.

**Table 1.** Visual test scores. Visual acuity scores are reported for high contrast (HC) and low contrast (LC) charts presented at the 63 cm viewing distance, and expressed as raw visual acuity scores (VAS). The corresponding logMAR scores are presented below in italics. Square brackets indicate the minimum and maximum scores across participants in each age group. Contrast sensitivity (CS) scores for young and older participants correspond to median log units across all participants in each age group.

|  | HC 63 | LC 63 | CS |
| --- | --- | --- | --- |
| Young | 108 [95, 110]<br>*-0.16 [0.10, -0.20]* | 99 [94, 104]<br>*0.02 [0.12, -0.08]* | 1.95 [1.95, 2.25] |
| Older | 98 [93, 105]<br>*0.04 [0.14, -0.10]* | 89 [82, 95]<br>*0.22 [0.36, 0.10]* | 1.95 [1.65, 1.95] |

The study was approved by the local ethics committee at the College of Science and Engineering, University of Glasgow (approval no. FIMS00740), and conducted in line with the British Psychological Society ethics guidelines. Informed written consent was obtained from each participant before the study. Participants were compensated £6/h.

### Stimuli

We used a set of 10 grey-scaled, front-view photographs of faces, oval cropped to remove external features, and pasted onto a uniform grey background (36). The pictures spanned 9.3° × 9.3° of visual angle; the face oval was 4.9° × 7.0° of visual angle. A unique image was presented on each trial by introducing phase noise (70% phase coherence) into the face images (37). Noisy textures were created by fully randomising the phase of the face images (0% phase coherence). All stimuli had the same amplitude spectrum, set to the mean amplitude of the face images. Face and noise images were revealed through ‘bubble masks’, i.e. masks containing 10 two-dimensional Gaussian apertures (sigma = 0.36°), with the constraint that the center of the aperture remained in the face oval (23). We wrote our experiments in MATLAB using the Psychophysics Toolbox extensions (38–40).

### Procedure

Participants came in for two experimental sessions on separate days. During each session, we asked participants to minimise movement and blinking, or blink only when hitting a response button. We maintained a viewing distance of 80 cm using a chinrest.

In each experimental session, participants completed 12 blocks of 100 trials each, seated in a sound-attenuated booth. The first block was a practice block of images without bubble masks. As such, across the two sessions participants performed 200 trials without bubble masks, and 2,200 trials with bubble masks. Practice blocks used a set of 10 face identities and 10 unique noise textures, each repeated 5 times were randomized within each block. Each practice session lasted about 60 to 75 minutes, including breaks, but excluding EEG electrode application.

On each trial, we instructed participants to categorise stimuli as fast and as accurately as possible, by pressing ‘1’ for face, and ‘2’ for texture on the numerical pad of a keyboard, using the index and middle finger of their dominant hand. Each trial began with a small black fixation cross (12 × 12 pixels, 0.4° × 0.4° of visual angle) displayed at the centre of the monitor screen for a random time interval of 500 to 1000 ms, followed by an image of a face or a texture presented for 7 frames (∼82 ms). After the stimulus, a blank grey screen was displayed until the participant responded. The fixation cross, the stimulus and the blank response screen were all displayed on a uniform grey background with mean luminance of ∼43 cd/m^2^. After each block of 100 trials, participants could take a break, and they received feedback on their performance in the previous block and on their overall performance in the experiment (median reaction time and percentage of correct responses). The next block started after participants pressed a key.

### EEG recording and pre-processing

We recorded EEG data at 512 Hz using a 128-channel Biosemi Active Two EEG system (Biosemi, Amsterdam, the Netherlands). Four additional UltraFlat Active Biosemi electrodes were placed below and at the outer canthi of both eyes. Electrode offsets were kept between ±20 µV.

EEG data were pre-processed using MATLAB 2013b and the open-source EEGLAB toolbox (41, 42). Data were first average-referenced and de-trended. Two types of filtering were then performed. First, data were band-pass filtered between 1 Hz and 30 Hz using a non-causal fourth order Butterworth filter. Independently, another dataset was created in which data were pre-processed with fourth order Butterworth filters: a high-pass causal filter at 2 Hz and a low-pass non-causal filter at 30 Hz, to preserve accurate timing of onsets (43–46).

Data from both datasets were then downsampled to 500 Hz, and epoched between − 300 and 1000 ms around stimulus onset. Mean baseline was removed from the causal-filtered data, and channel mean was removed from each channel in the non-causal-filtered data in order to increase the reliability of Independent Component Analysis (ICA) (47). Noisy electrodes and trials were then detected by visual inspection of the non-causal dataset, and rejected on a participant-by-participant basis. Following visual inspection, one young participant and one older participant were excluded from further analyses due to noisy EEG signal. Mutual Information (MI) analysis confirmed the lack of sensitivity to any facial features in these participants. The resulting sample size was 17 young and 18 older participants. In this sample, more noisy channels were on average removed from older than from young participants’ datasets (older participants: median = 10, min = 0, max = 24; young participants: median = 5, min = 0, max = 28; median difference = 4 [2, 7]). More noisy Bubble trials were also removed from older than from young participants’ datasets (trials included in analyses, older participants: median 2130, min 1987, max 2180; young participants: median 2178, min 2023, max 2198; median difference = 42 [23, 64]).

Subsequently, we performed ICA on the non-causal filtered dataset using the Infomax algorithm as implemented in the *runica* function in EEGLAB (42, 48). The ICA weights were then applied to the causal filtered dataset to ensure removal of the same components, and artifactual components were rejected from both datasets (median = 4, min = 1, max = 27 for one older participant who displayed excessive blink activity; the second max was 17). Then, baseline correction was performed again, and data epochs were removed based on an absolute threshold value larger than 100 µV and the presence of a linear trend with an absolute slope larger than 75 µV per epoch and R^2^ larger than 0.3. The median number of bubble trials accepted for analysis was, out of 1100, for older participants: face trials = 1069 [min=999, max=1092]; noise trials = 1067 [min=986, max=1088]; for young participants: face trials = 1090 [min=1006, max=1100]; noise trials = 1089 [min=1014, max=1098]. Finally, we computed single-trial spherical spline current source density waveforms using the CSD toolbox (49, 50). CSD waveforms were computed using parameters 50 iterations, m=4, lambda=10^-5^. The head radius was arbitrarily set to 10 cm, so that the ERP units are µV/cm^2^. The CSD transformation is a spatial high-pass filtering of the data, which sharpens ERP topographies and reduces the influence of volume-conducted activity. CSD waveforms are also reference-free.

### Electrode selection

We performed detailed analyses on the subset of four posterior midline electrodes that are sensitive to face features or conjunction of features: from top to bottom CPz, Pz, POz, Oz (23, 51). We only report the results of Oz because the other three electrodes had weak mutual information to face features in the two groups. We also selected two posterior-lateral electrodes, one in the right hemisphere (right electrode, RE), and one in the left hemisphere (left electrode, LE) by measuring the difference between all bubble face trials and all bubble noise trials at all posterior-lateral electrodes, squaring it, and selecting the left and the right electrodes that showed the maximum difference in the period 130-250 ms. Across participants, the selected LE and RE were P7/8, or PO7/8, or their immediate neighbours. These electrodes are typically associated with large face ERPs in the literature.

### Event-related potentials

We compared the amplitude and latency of the N170 between the two age groups. To this end, we computed mean ERPs across trials for each participant, separately for face and noise trials, and for practice (without Bubbles) and regular (with Bubbles) trials. For ERPs recorded at the lateral-occipital electrode in the right hemisphere (RE), we defined the N170 peak in individual participants as the minimum mean ERP between 110-230 ms, and considered separately its latency and amplitude.

### Statistical analyses

We conducted statistical analyses using Matlab 2013b and the LIMO EEG toolbox (52). Throughout the paper, square brackets indicate 95% confidence intervals computed using the percentile bootstrap technique, with 1,000 bootstrap samples. Unless otherwise stated, median values are Harrell-Davis estimates of the 2^nd^ quartile (53).

### Mutual Information

We used mutual information (MI) to quantify the dependence between stimulus features and behavioural and brain responses (9,51,54–56). We binned pixel visibility across trials (due to random bubbles sampling), and behavioural and EEG responses into three equiprobable bins (for details, see (23)). We then calculated several MI quantities in single participants: MI(PIX; RT) to establish the relationship between image pixels and reaction times; MI(PIX; CORRECT) to establish the relationship between image pixels and correct responses; MI(PIX; RESP) between pixels and response category; and MI(PIX; ERP) to establish the relationship between image pixels and ERPs. We computed these quantities separately for face and noise trials. To control for the variable number of trials in each participant arising as a result of EEG pre-processing, we scaled every MI quantity for every participant by a factor of *2Nln2* (57), using the formula:

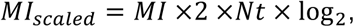

where *MI* refers to mutual information values, and *Nt* is the number of trials. *MI*_*scaled*_, therefore, reflects a measure of MI adjusted for a systematic upward bias in the information estimate that might arise due to limited data sampling, especially if the numbers of trials in the two age groups are systematically different. It also converts MI to be the effect size for a log-likelihood test of independence (58). All group-difference analyses were performed using the scaled MI values.

### Mutual Information: Classification images

We refer to MI between pixels and behaviour or ERPs as classification images: they reveal the image pixels associated with modulations of the responses. We computed the MI(PIX, ERP) classification images at every time point within the first 400 ms following stimulus onset, using the non-causal and causal-filtered datasets, and at each of the 6 electrodes specified above. We summarized each classification image with its maximum MI and reported these time courses per electrode.

#### Single-subject analyses

To establish the statistical significance of the classification image pixels while controlling for multiple comparisons arising from testing at multiple pixels, we performed a permutation test coupled with the Threshold-Free Cluster Enhancement (TFCE) technique (59) on individual participants’ data (23).

#### Feature Redundancy

We computed co-information (coI) (60) (equivalent to interaction information (61) but with opposite sign for three variables) to quantify the triple dependence between eye visibility, reaction times and brain responses (9,62–65), i.e. coI(eye; RT; EEG). Positive values of coI quantify redundancy, specifically here redundant or overlapping information about the stimulus eye visibility, which is common to both EEG and reaction time responses. We computed co-Information redundancy with the following expression:

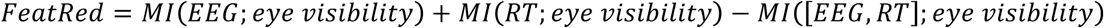

We used here Gaussian-Copula Mutual Information (GCMI) a semi-parametric rank based estimator suitable for continuous variables (9) (see also *Supplementary Methods*) and considered a two-dimensional EEG signal consisting of voltage together with its instantaneous temporal derivative (see *Supplementary Methods*).

If there is less information about eye visibility available from EEG and RT considered jointly, MI([EEG RT]; eye visibility), than there is when the two responses are considered independently, MI(EEG; eye visibility)+MI(RT; eye visibility), then this shows that part of the relationship quantified by MI(EEG; eye visibility) overlaps with that quantified by MI(RT; eye visibility). As such, redundancy quantifies the overlapping information content (i.e. a common driving effect) within both EEG and reaction time, about eye visibility. We computed this measure at all time points and electrodes for each participant independently. Then, for each participant, we and plotted the time course of redundancy at occipital-temporal electrodes specified above (i.e. RE and LE) between 0 and 400 ms post-stimulus.

### Group Redundancy

We computed group-level redundancy to quantify the degree to which median RT and individual peak redundancy time provide a common prediction of the age group (young vs. older) across individual participants. For these analyses, we quantized RT and peak redundancy time into three equally populated bins (splitting on tertiles of the distribution across participants) and calculated MI from the standard discrete definition (9, 66). We applied Miller-Madow bias correction to the individual MI estimated before calculating group redundancy as:

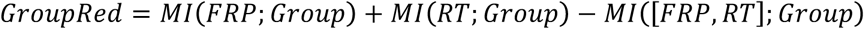

where FRP is Feature Redundancy Peak time, RT is reaction time and Group is age group for each participant. We determined bootstrap confidence intervals via resampling participants with replacement 1,000 times. Since group redundancy is bounded above by the minimum of MI(FRP; Group) and MI(RT; Group), we normalized by that minimum to obtain redundancy as a percentage of the theoretical maximum (the smaller of the two pairwise MI’s).

### ERP onset analyses

We quantified ERP onsets using the causal-filtered datasets. To control for multiple comparisons, we used a bootstrap temporal clustering technique as implemented in LIMO EEG (52, 67).

#### ERP_STD_ onset

To test whether age-related delays reflect differences in the onset of afferent activity to the visual cortex, we looked at the time course of the standard deviation across electrodes of the mean ERP (ERP_STD_). ERP_STD_ provides a compact description of the global ERP response, summarizing each participant’s evoked brain activity across electrodes in one vector. This analysis was based on the notion that early visual activity can be characterized by a sudden increase in standard deviation of the mean ERP across electrodes (8). We computed the ERP_STD_ time course for each individual participant and mean baseline centered it. Then, we localised the first peak the minimum height of which was five times the height of any peak in the baseline. Then, using ARESLab toolbox (68), we built a piecewise-linear regression model with three basis functions using the Multivariate Adaptive Regression Splines (MARS) (69) method. This approach divided the data into two segments and fitted each segments with separate models (regression splines). Onsets were defined as the location in time of the first knot of the fitted spline, i.e. the point where division between the two models occurred.

#### MI onset

We quantified MI onsets using the same technique as with ERP_STD_ onsets.

### Topographic analyses

We computed topographic maps for each participant from the whole-scalp MI(PIX; ERP) results, and for the whole-scalp ERP results, at the individual MI peak, or ERP peak latency, respectively. Individual ERP topographic maps were squared. All topographic maps were interpolated and rendered in a 67 × 67 pixel image using the EEGLAB function *topoplot*, and then averaged across participants in each age group. Using the interpolated head maps, we then computed a hemispheric lateralisation index for each participant using MI results. First, we normalised MI values in each participant between 0 and 1. Then, we saved the maximum pixel intensity in the left and the right hemisphere (lower left and right quadrants of the interpolated image), excluding the midline. Finally, we computed the lateralisation index in each group as the ratio (MI_left_ - MI_right_) / (MI_left_ + MI_right_). Whole-scalp MI was strongest at posterior-lateral electrodes, and tended to be right lateralised in both groups (lateralisation index for face trials, young = −0.18 [−0.31, −0.05]; older = −0.23 [−0.37, − 0.09]; group difference = 0.07 [−0.07, 0.21]).

## Acknowledgments

This work has been funded by the UK Biotechnology and Biological Sciences Research Council (BBSRC) grant BB/J018929/1 awarded to NJVR, GAR, and PGS. KJ was supported by the BBSRC Doctoral Training Partnership (WestBio) Studentship. PGS received support from the Wellcome Trust (Senior Investigator Award, UK; 107802) and the Multidisciplinary University Research Initiative/Engineering and Physical Sciences Research Council (USA, UK; 172046-01).

## Data Availability

A reproducibility package with data and code will be available online as soon as possible.

## References

1. Salthouse TA. Aging and measures of processing speed. Biol Psychol. 2000 Oct;54(1–3):35–54.

2. Rousselet GA, Husk JS, Pernet CR, Gaspar CM, Bennett PJ, Sekuler AB. Age-related delay in information accrual for faces: evidence from a parametric, single-trial EEG approach. BMC Neurosci. 2009;10:114.

3. Rousselet GA, Gaspar CM, Pernet CR, Husk JS, Bennett PJ, Sekuler AB. Healthy aging delays scalp EEG sensitivity to noise in a face discrimination task. Front Psychol. 2010;1(JUL):1–14.

4. Gazzaley A, Clapp W, Kelley J, McEvoy K, Knight RT, D’Esposito M. Age-related top-down suppression deficit in the early stages of cortical visual memory processing. Proc Natl Acad Sci. 2008;105(35):13122–6.

5. Nakamura A, Yamada T, Abe Y, Nakamura K, Sato N, Horibe K, et al. Age-related changes in brain neuromagnetic responses to face perception in humans. Vol. 312, Neuroscience Letters. 2001.

6. Wiese H, Schweinberger SR, Hansen K. The age of the beholder: ERP evidence of an own-age bias in face memory. Neuropsychologia. 2008;46(12):2973–85.

7. Gosselin F, Schyns PG. Bubbles: A technique to reveal the use of information in recognition tasks. Vision Res. 2001;41(17):2261–71.

8. Foxe JJ, Simpson G V. Flow of activation from V1 to frontal cortex in humans: A framework for defining “early” visual processing. Exp Brain Res. 2002;142(1):139–50.

9. Ince RAA, Giordano BL, Kayser C, Rousselet GA, Gross J, Schyns PG. A statistical framework for neuroimaging data analysis based on mutual information estimated via a gaussian copula. Hum Brain Mapp. 2017 Mar;38(3):1541–73.

10. Chaby L, Narme P, George N. Older adults’ configural processing of faces: Role of second-order information. Psychol Aging. 2011 Mar;26(1):71–9.

11. Obermeyer S, Kolling T, Schaich A, Knopf M. Differences between old and young adults’ ability to recognize human faces underlie processing of horizontal information. Front Aging Neurosci. 2012;4(APR):1–9.

12. van Rijsbergen NJ, Jaworska K, Rousselet GA, Schyns PG. With age comes representational wisdom in social signals. Curr Biol. 2014 Dec 1;24(23):2792–6.

13. Lott LA, Haegerstrom-Portnoy G, Schneck ME, Brabyn JA. Face recognition in the elderly. Optom Vis Sci. 2005;82(10):874–81.

14. Owsley C, Sekuler R, Boldt C. Aging and low-contrast vision: face perception. Investig Ophthalmol Vis Sci. 1981;21(2):362–5.

15. Creighton SE, Bennett PJ, Sekuler AB. Classification images characterize age-related deficits in face discrimination. Vision Res. 2018 Aug 1;

16. Éthier-Majcher C, Joubert S, Gosselin F. Reverse correlating trustworthy faces in young and older adults. Front Psychol. 2013 Sep 5;4:592.

17. Sergent J, Ohta S, MacDonald B. Functional neuroanatomy of face and object processing. A positron emission tomography study. Brain. 1992 Feb;115 Pt 1:15–36.

18. Bieniek MM, Frei LS, Rousselet GA. Early ERPs to faces: Aging, luminance, and individual differences. Front Psychol. 2013;4(MAY).

19. Bieniek MM, Bennett PJ, Sekuler AB, Rousselet GA. A robust and representative lower bound on object processing speed in humans. Foxe J, editor. Eur J Neurosci. 2016 Jul 1;44(2):1804–14.

20. Price D, Tyler LK, Neto Henriques R, Campbell KL, Williams N, Treder MS, et al. Age-related delay in visual and auditory evoked responses is mediated by white- and grey-matter differences. Nat Commun. 2017 Jun 9;8:15671.

21. Schyns PG, Petro LS, Smith ML. Dynamics of Visual Information Integration in the Brain for Categorizing Facial Expressions. Curr Biol. 2007;17(18):1580–5.

22. Smith ML, Gosselin F, Schyns PG. Receptive fields for flexible face categorizations. Psychol Sci. 2004;15(11):753–61.

23. Rousselet GA, Ince RAA, van Rijsbergen NJ, Schyns PG. Eye coding mechanisms in early human face event-related potentials. 2014;14:1–24.

24. Zanto TP, Toy B, Gazzaley A. Delays in neural processing during working memory encoding in normal aging. Neuropsychologia. 2010 Jan;48(1):13–25.

25. Bentin S, Allison T, Puce A, Perez E, McCarthy G. Electrophysiological studies of face perception in humans. J Cogn Neurosci. 1996;8(6):551–65.

26. Itier RJ, Alain C, Sedore K, McIntosh AR. Early face processing specificity: it’s in the eyes! J Cogn Neurosci. 2007;19(11):1815–26.

27. Van Rijsbergen NJ, Schyns PG. Dynamics of trimming the content of face representations for categorization in the brain. PLoS Comput Biol. 2009;5(11).

28. Kolev V, Falkenstein M, Yordanova J. Motor-response generation as a source of aging-related behavioural slowing in choice-reaction tasks. Neurobiol Aging. 2006 Nov;27(11):1719–30.

29. Yordanova J, Kolev V, Hohnsbein J, Falkenstein M. Sensorimotor slowing with ageing is mediated by a functional dysregulation of motor-generation processes: evidence from high-resolution event-related potentials. Brain. 2004 Feb 1;127(2):351–62.

30. Tang H, Buia C, Madhavan R, Crone NE, Madsen JR, Anderson WS, et al. Spatiotemporal Dynamics Underlying Object Completion in Human Ventral Visual Cortex. Neuron. 2014;83(3):736–48.

31. Sekuler AB, Gold JM, Murray RF, Bennett P. Visual completion of partly occluded objects: insights from behavioral studies. Neuro-ophtalmology. 2000;23:165–8.

32. Wang Y, Zhou Y, Ma Y, Leventhal AG. Degradation of Signal Timing in Cortical Areas V1 and V2 of Senescent Monkeys. Cereb Cortex. 2005 Apr 1;15(4):403–8.

33. Raz N, Lindenberger U, Rodrigue KM, Kennedy KM, Head D, Williamson A, et al. Regional Brain Changes in Aging Healthy Adults: General Trends, Individual Differences and Modifiers. Cereb Cortex. 2005 Nov 1;15(11):1676–89.

34. Peters A. The effects of normal aging on myelinated nerve fibers in monkey central nervous system. Front Neuroanat. 2009 Jul 6;3:11.

35. Elliott DB, Sanderson K, Conkey A. The reliability of the Pelli-Robson contrast sensitivity chart. Ophthalmic Physiol Opt. 1990 Jan;10(1):21–4.

36. Gold J, Bennett PJ, Sekuler AB. Identification of band-pass filtered letters and faces by human and ideal observers. Vision Res. 1999;39(21):3537–60.

37. Rousselet GA, Pernet CR, Bennett PJ, Sekuler AB. Parametric study of EEG sensitivity to phase noise during face processing. BMC Neurosci. 2008;9:98.

38. Brainard DH. The Psychophysics Toolbox. Spat Vis. 1997;10(4):433–6.

39. Kleiner M, Brainard DH, Pelli D. What’s new in Psychtoolbox-3? Perception. 2007;36(ECVP Abstract Supplement).

40. Pelli DG. The VideoToolbox software for visual psychophysics: transforming numbers into movies. Spat Vis. 1997;10(4):437–42.

41. Delorme A, Mullen T, Kothe C, Akalin Acar Z, Bigdely-Shamlo N, Vankov A, et al. EEGLAB, SIFT, NFT, BCILAB, and ERICA: new tools for advanced EEG processing. Comput Intell Neurosci. 2011;2011:130714.

42. Delorme A, Makeig S. EEGLAB: an open source toolbox for analysis of single-trial EEG dynamics including independent component analysis. J Neurosci Methods. 2004;134:9–21.

43. Acunzo DJ, MacKenzie G, van Rossum MCW. Systematic biases in early ERP and ERF components as a result of high-pass filtering. J Neurosci Methods. 2012;209(1):212–8.

44. Luck SJ. An introduction to the event-related potential technique. MIT Press; 2005. 374 p.

45. Rousselet GA. Does Filtering Preclude Us from Studying ERP Time-Courses? Front Psychol. 2012;3:131.

46. Widmann A, Schröger E. Filter effects and filter artifacts in the analysis of electrophysiological data. Front Psychol. 2012;3:233.

47. Groppe DM, Makeig S, Kutas M. Identifying reliable independent components via split-half comparisons. Neuroimage. 2009 May;45(4):1199–211.

48. Delorme A, Sejnowski T, Makeig S. Enhanced detection of artifacts in EEG data using higher-order statistics and independent component analysis. Neuroimage. 2007 Feb;34(4):1443–9.

49. Kayser J. Current source density (CSD) interpolation using spherical splines - CSD Toolbox (Version 1.1). New York State Psychiatric Institute: Division of Cognitive Neuroscience; 2009.

50. Tenke CE, Kayser J. Generator localization by current source density (CSD): implications of volume conduction and field closure at intracranial and scalp resolutions. Clin Neurophysiol. 2012 Dec;123(12):2328–45.

51. Schyns PG, Thut G, Gross J. Cracking the code of oscillatory activity. PLoS Biol. 2011;9(5).

52. Pernet CR, Chauveau N, Gaspar C, Rousselet GA, Pernet CR, Chauveau N, et al. LIMO EEG: a toolbox for hierarchical LInear MOdeling of ElectroEncephaloGraphic data. Comput Intell Neurosci. 2011;2011:831409.

53. Harrell FE, Davis CE. A new distribution-free quantile estimator. Biometrika. 1982;69(3):635–40.

54. Ince RAA, Petersen RS, Swan DC, Panzeri S. Python for information theoretic analysis of neural data. Front Neuroinform. 2009;3:4.

55. Park H, Ince RAA, Schyns PG, Thut G, Gross J. Frontal Top-Down Signals Increase Coupling of Auditory Low-Frequency Oscillations to Continuous Speech in Human Listeners. Vol. 25, Current Biology. 2015.

56. Kayser SJ, Ince RAA, Gross J, Kayser C. Irregular Speech Rate Dissociates Auditory Cortical Entrainment, Evoked Responses, and Frontal Alpha. J Neurosci. 2015;35(44).

57. Ince RAA, Mazzoni A, Bartels A, Logothetis NK, Panzeri S. A novel test to determine the significance of neural selectivity to single and multiple potentially correlated stimulus features. J Neurosci Methods. 2012;210(1):49–65.

58. Sokal RR, Rohlf FJ. Biometry?: the principles and practice of statistics in biological research. 4th Editio. W.H. Freeman; 2012.

59. Smith SM, Nichols TE. Threshold-free cluster enhancement: Addressing problems of smoothing, threshold dependence and localisation in cluster inference. Neuroimage. 2009;44(1):83–98.

60. Bell AJ. The co-information lattice. In: Proceedings of the 4th International Symposium on Independent Component Analysis and Blind Signal Separation (ICA2003), Nara, Japan, 1-4 April. 2003. p. 921–6.

61. McGill WJ. Multivariate information transmission. Psychometrika. 1954 Jun;19(2):97– 116.

62. Ince RAA. Measuring Multivariate Redundant Information with Pointwise Common Change in Surprisal. Entropy. 2017 Jun 29;19(7):318.

63. Zhan J, Ince RAA, Van Rijsbergen NJ, Schyns PG. Dynamic construction of reduced representations in the brain for perceptual decision behavior. Curr Biol (in Press. 2018 Nov 20;

64. Ince RAA, van Rijsbergen NJ, Thut G, Rousselet GA, Gross J, Panzeri S, et al. Tracing the Flow of Perceptual Features in an Algorithmic Brain Network. Sci Rep. 2015 Nov 4;5(1):17681.

65. Ince RAA, Jaworska K, Gross J, Panzeri S, van Rijsbergen NJ, Rousselet GA, et al. The Deceptively Simple N170 Reflects Network Information Processing Mechanisms Involving Visual Feature Coding and Transfer Across Hemispheres. Cereb Cortex. 2016 Oct 17;26(11):4123–35.

66. Cover TM, Thomas JA. Elements of Information Theory. Elements of Information Theory. 2005. 1–748 p.

67. Pernet CR, Latinus M, Nichols TE, Rousselet GA. Cluster-based computational methods for mass univariate analyses of event-related brain potentials/fields: A simulation study. J Neurosci Methods. 2015;250:85–93.

68. Jekabsons G. ARESLab: Adaptive Regression Splines toolbox for Matlab/Octave. 2015.

69. Friedman JH. Multivariate Adaptive Regression Splines. Ann Stat. 1991 Mar;19(1):1– 67.

