## Supplementary Material for "Healthy ageing delays the neural processing of face features relevant for behaviour by 40 ms"

### Supplementary Methods

#### Reverse Analysis

To quantify how the presence of the eyes modulated behavioural and brain responses, we ran a reverse analysis (1,2). First, we created the eye mask by centring a circle (radius = 15 pixels) on the pixel that showed the maximum MI value in the group-averaged classification image (for brain MI(PIX, ERP) information), separately for the left and for the right eye. We then summed pixel values revealed through single-trial Bubble masks that fell within the boundaries of each eye mask independently, and within both eye masks together, to provide an estimate of eye region visibility. We then split these values into ten equally populated bins ranging from the lowest to the highest sum values and computed the median RT and the mean percent correct for each bin. Next, we quantified the effect of eye visibility on behavioural judgments by calculating the RT and percent correct difference between the 10$^{th}$ and the 1$^{st}$ bin. We then repeated this analysis with single-trial ERP distributions: we averaged the ERPs corresponding to each bin, separately for the left and right lateral electrodes. We then computed the N170 amplitude and latency in every participant and for each eye mask, for the lowest (1$^{st}$ bin) and the highest (10$^{th}$ bin) sum values, separately for the left and right electrodes. Given that the N170 on Bubble trials was delayed with respect to that on practice trials in both groups, we defined the N170 as the minimum in the time window 150 to 250 ms following stimulus onset in ERPs low-pass filtered at 20 Hz using a fourth order Butterworth non-causal filter. We then computed the differences between high and low amplitude and latency values for each group separately.

#### Measures of effect size

We estimated the size of the between-group differences using two robust techniques: Cliff's delta and the median of all pairwise differences. Cliff's *delta* (3,4) is related to the Wilcoxon-Mann-Whitney *U* statistic and estimates the probability that a randomly selected observation from one group is larger than a randomly selected observation from another group, minus the reverse probability. Cliff's *delta* ranges from 1 when all values from one group are higher than the values from the other group, to -1 when the reverse is true. Completely overlapping distributions have a Cliff's *delta* of 0. In line with Cliff's delta approach, we also calculated all pairwise differences between young and older participants on the measures of interest (reaction times, percent corrects, N170 latencies and amplitudes), and took the median of the distribution of these differences. This way of measuring effect sizes enabled us to provide information about the typical difference between any two observations from two groups (5).

#### Comparison of MI values obtained with different methods

In the main manuscript, we used an approach where data were quantized into bins, and MI was estimated over the resulting discrete spaces. However, the binning approach can be sensitive to the problem of limited sampling bias. To test the results described in the main manuscript, here we computed a new estimator of MI that can be used with continuous variables: Gaussian-Copula Mutual Information (GCMI) (6). GCMI is a semi-parametric estimator that utilizes the concept of copulas (7), statistical structures that express the relationship between two random variables, independently of their marginal distributions. The copula method does not require the binning step and overcomes this problem while being robust and computationally efficient (6).

In single participants, we calculated MI between visibility of the left eye (a scalar value obtained as a sum of pixel visibility within the circular left eye mask on each trial, cf. main manuscript) and EEG voltage over the period of -300 ms before to 1000 ms after stimulus onset (*regular* MI, Supplementary Table S1). We also computed the temporal gradient of the EEG voltage (dEEG) on each trial to account for the temporal relationship between neighbouring time points, and then combined the EEG voltage and its temporal gradient into a bivariate response (*gradient* MI, Supplementary Table S1). We then calculated the time course of MI about the eye visibility in the bivariate response: MI(eye, [EEG dEEG]). Considering the gradient response together with the voltage smoothes out the artifactual dips in MI time courses, occurring at time points of zero-crossings when EEG voltages change sign. It also introduces information about the shape of the ERP, otherwise missing from just considering instantaneous amplitudes. As such, the bivariate time course provides a clearer picture of the intervals over which stimulus variability modulates the EEG signal (6). Finally, to check whether information about the eye visibility was coded in a more distributed manner across the scalp in older compared with young participants, we ran Principal Component Analysis (PCA) across electrodes at each time point in individual participants. Then, we computed MI between the top four components and eye visibility (*multivariate* MI, Supplementary Table S1).

All analyses yielded group differences comparable to those obtained with the binning method and described in the main manuscript: MI about the left eye was delayed by about 37 - 47 ms in older compared with young participants, and peak MI in older participants was 49 - 59% the size of that in young participants. As such, the binning method was sensitive enough to provide a good description of age-related differences in eye coding. Furthermore, analyses restricted to a single lateral-occipital electrode in each hemisphere (see main manuscript) were sufficient to describe age-related differences in processing of facial information in our study.

Finally, to check whether time courses of max MI (see *Results*, main manuscript) provided a good description of the data, we also computed the *sum* of MI across pixels instead of the *maximum,* and computed its peak MI latency and amplitude, along with the group differences (see Supplementary Table S2). By considering the sum, we noticed that the older to young ratio of MI increased to about 74% [61, 90] at LE, and 80% [64, 95] at RE – an effect likely driven by more distributed MI in older participants (see Supplementary Figure S10), albeit not significant at single participant level (cf. Figure 4, main manuscript).

### Supplementary Results

#### Presence of the eyes was crucial for older participants to detect a face

Young participants detected faces more accurately than older participants (bubble trials, mean young = 93% [92, 94] vs. older = 82% [80, 85]; group difference = 11 percentage points (PP) [8, 14]), although both groups reached 98% [97, 98] on practice trials in which face and noise images were presented without bubbles. Older adults found the Bubbles task more challenging (bubble-practice difference, young = -4 PP [-6, -3] vs. older = -15 PP [-18, -11]; group difference = -11 PP [-14, -7]). Specifically, older participants had higher numbers of noise responses on face trials, while both groups were similarly accurate on noise trials (face trials, young = 91% correct vs. older = 75% correct; group difference = 15 PP [8, 19]; noise trials, young = 95% vs. older = 92%; group difference = 4 PP [1, 13]; group difference of face-noise difference = 7 PP [0, 15]).

Lower accuracy on bubble trials is in line with other studies showing age-related deterioration in performance on tasks involving perception of partially occluded objects. For example, older participants were less accurate and needed more stimulus information in tasks requiring perceptual closure (8–10), perceptual organization (11), contour integration (12) or perception of incomplete/fragmented figures (13–15). The underlying causes of such perceptual difficulties remain unclear, although it has been suggested that they arise as a consequence of heightened noise or variability associated with internal stimulus representation in the neural system (16), or as a result of a deficit in inhibitory control of interfering/irrelevant information (15).

To understand the face information associated with behaviour, we used Mutual Information (MI; see *Materials and Methods*) to compute the relationship between pixel visibility and correct vs. incorrect responses. Eye region pixels were strongly associated with correct responses in only a few young participants and almost all older participants, suggesting that young participants used any feature to do the task, in contrast to older participants who needed to see the eye region to correctly detect a face (N young = 4/17, N older = 16/18, Supplementary Figure S1 A-B). This reliance on the eyes was confirmed in the average classification image of the difference between the groups (Supplementary Figure S1 C).

We confirmed these results with a reverse analysis (see *Supplementary Methods*) that showed, in young adults, a 53 ms RT gain [45, 61] and 15 percentage points (PP) accuracy gain [10, 20] when the left eye was visible – vs. 91 ms [68, 114] and

37 PP [25, 48] gains in older participants (Supplementary Figure S2). For the right eye, these gains were respectively 17 ms [10, 24] and 7 PP [4, 11] in young participants; and 16 ms [2, 30] and 28 PP [19, 36] in older participants (see Supplementary Table S3 for effect sizes of group differences).

To demonstrate a difference in behavioural strategy between the groups, Supplementary Figure S3 (bottom panel) shows that on trials without any eye visibility (either left or right), young participants could still detect faces accurately, whereas older participants could not (bin 1, young: min = 56%, median = 72%, max = 92%; older: min = 9%, median = 31%, max = 94%; see also Supplementary Figures S4 – S6 for reaction times, and for noise trials).

In sum, our behavioural results reveal that all participants used the eyes to detect faces from noise. However, whereas the eyes were all older participants could use to make correct responses, young participants could also use any other face feature; demonstrating a strategy difference in older participants.

#### Effect sizes for brain sensitivity to image structure

To compare the EEG sensitivity to faces across the two age groups, we computed Cliff's *delta* estimates for the difference between ERPs to face and noise practice trials at every time point and electrode of every participant. Then, we saved the maximum *delta* value across time points and electrodes in each participant, and computed the median across participants in each age group. The EEG face sensitivity to full images was similar across the two groups (young-older difference 0.01 [-0.07, 0.09]; median Cliff's *delta* in young participants = 0.64 [0.58, 0.68], median Cliff's *delta* in older participants = 0.60 [0.54, 0.69]).

#### N170 Latency and amplitude code the presence of the eyes differentially in young and older adults

To uncover the functional role of the N170 in coding task-relevant information in older adults, using a reverse analysis we sought to directly investigate how eye processing related to the N170 latency and amplitude (Supplementary Figure S9; see *Supplementary Methods;* see also (1,2)). As shown below (see Supplementary Figure S9), we found that pixels in the contralateral eye region modulate amplitude and latency distributions of single-trial N170 in young participants (2), but mostly amplitude distributions in older participants.

Specifically, the N170 reconstructed from trials with high contralateral eye visibility on RE preceded and was larger than the N170 reconstructed from trials with low contralateral eye visibility (latency effect, young = 24 ms [17, 31] vs. older = 5 ms [-2, 11]; for amplitude effects, see Supplementary Table S5). This latency effect was weaker at LE (young = 12 ms [8, 17]; older = 1 ms [-6, 9]), stronger in young than in older participants (for effect size estimates, see Table S4), and stronger at RE than LE in young compared to older participants (young = -7 ms [-12, -3] vs. older = -3 ms [-7, 3]; group difference = -5 ms [-15, 0]). Surprisingly, contralateral eye amplitude modulation at LE was *weaker* in young than in older participants, while similar across groups at RE (LE, young = 153% [140, 166]; older = 191% [157, 225]; see also Supplementary Tables S4 and S5).

The presence of the ipsilateral eye had opposite effects on the N170 latency in the two groups: high ipsilateral eye visibility was associated with earlier N170 in young participants and later N170 in older participants (RE, young = 2 ms [1, 4] vs. older = -4 ms [-1, -8]; LE, young = 6 ms [2, 10] vs. older = -4 ms [-1, -8]).

In sum, our results suggest that ageing affects the N170 coding of the eye by showing that contra-lateral eye pixels modulate the amplitude and latency of the N170 in young participants (2), and mostly amplitude in older participants.

### Supplementary Tables

**Table S1** Supplementary MI results.

We computed continuous MI between eye visibility and EEG (***regular***), bivariate MI between eye visibility and combined EEG and its temporal gradient (***gradient***) and MI between eye visibility and PCA components (***multivariate***) in single subjects. Values correspond to the Harrell-Davis estimates of the median latency (expressed in milliseconds) of maximum MI, and maximum MI (expressed in bits) computed across young and older participants and at the left occipital-temporal (LE) and the right occipital-temporal sensors separately. Group differences are expressed as the median of all pairwise differences between participants in each group. Additionally, the group difference in MI amplitudes is expressed as the ratio of the median amplitude in older to the median amplitude in young participants. Square brackets indicate 95% confidence intervals.

| | Latency (ms) | | Amplitude (bits) | |
|---|---|---|---|---|
| ***regular*** | LE | RE | LE | RE |
| young | 165 [158, 177] | 162 [158, 172] | 0.07 [0.04, 0.11] | 0.12 [0.09, 0.14] |
| older | 212 [195, 249] | 201 [191, 218] | 0.04 [0.03, 0.06] | 0.05 [0.04, 0.08] |
| difference | 47 [29, 89] | 39 [25, 58] | 0.03 [0, 0.07] *ratio: 0.59 [0.39, 1.03]* | 0.06 [0.03, 0.09] *ratio: 0.51 [0.35, 0.77]* |
| ***gradient*** | | | | |
| young | 166 [157, 188] | 164 [156, 175] | 0.07 [0.05, 0.11] | 0.12 [0.09, 0.14] |
| older | 213 [196, 256] | 201 [189, 219] | 0.04 [0.03, 0.05] | 0.05 [0.04, 0.08] |
| difference | 47 [19, 84] | 37 [21, 56] | 0.03 [0, 0.07] *ratio: 0.58 [0.36, 1.04]* | 0.07 [0.03, 0.09] *ratio: 0.49 [0.33, 0.72]* |
| ***multivariate*** | | | | |
| young | 171 [166, 183] | | 0.09 [0.08, 0.12] | |
| older | 210 [195, 237] | | 0.06 [0.04, 0.08] | |
| difference | 38 [20, 70] | | 0.03 [0.01, 0.07] *ratio: 0.57 [0.39, 0.85]* | |

**Table S2** Sum of MI across pixels increases the ratio of older/young MI

Values correspond to the Harrell-Davis estimates of the median peak latency (expressed in milliseconds) of the time courses of MI sum across pixels, and peak amplitude (expressed in bits) computed across young and older participants and at the left occipital-temporal (LE) and the right occipital-temporal sensors separately. Group differences are expressed as the median of all pairwise differences between participants in each group. Additionally, the group difference in MI amplitudes is expressed as the ratio of the median amplitude in older to the median amplitude in young participants. Square brackets indicate 95% confidence intervals.

| | Latency (ms) | | Amplitude | |
|---|---|---|---|---|
| ***sum*** | LE | RE | LE | RE |
| young | 171 [162, 178] | 165 [162, 169] | 8 [6.7, 9.8] | 9.5 [7.9, 10.8] |
| older | 216 [194, 234] | 206 [194, 232] | 5.5 [5.1, 6.7] | 7 [6, 8.8] |
| difference | 45 [21, 70] | 41 [28, 67] | 2.5 [0.1, 4.5] | 2.5 [0.3, 4.2] |
| | | | *Ratio $_{older/young}$ (%)* | |
| | | | *0.74 [0.61, 0.90]* | *0.80 [0.64, 0.95]* |

**Table S3** Effect size estimates for group differences in high vs. low eye visibility.

Values correspond to group differences between high vs. low eye visibility differences in RT (expressed in milliseconds) and percent correct (percentage points). Values correspond to the median of all pairwise differences between each young and every older participant. Negative accuracy and RT differences indicate that older participants were more affected by eye visibility than young participants. A corresponding Cliff's *delta* is shown in italics. Square brackets indicate 95% confidence intervals.

| | Accuracy scores | | Reaction times | |
|---|---|---|---|---|
| | Face trials | Noise trials | Face trials | Noise trials |
| Left eye | -21.3 [-30.5, -10.3] | 1.9 [-0.3, 3.8] | 40.6 [19, 60.3] | 4.9 [-7.4, 13.2] |
| | *-0.71 [-0.93, -0.44]* | *0.37 [-0.02, 0.70]* | *0.71 [0.41, 0.94]* | *0.16 [-0.23, 0.54]* |
| Right eye | -19.2 [-26, -10.1] | 4.0 [0.9, 6.7] | -2.5 [-16.2, 9.8] | 11.2 [-7.9, 27.2] |
| | *-0.73 [-0.94, -0.44]* | *0.50 [0.12, 0.82]* | *-0.08 [-0.49, 0.31]* | *0.25 [-0.17, 0.62]* |

**Table S4** Effect size estimates for N170 eye coding.

Effect size estimates for group differences (young-older) in N170 latency (LAT) and amplitude (AMP) for different facial features visibility, at left (LE) and right electrodes (RE). Values correspond to differences in median latencies in milliseconds, and median amplitudes in percentage points. Square brackets indicate 95% confidence intervals. A corresponding Cliff's delta estimate is shown in italics.

| | N170 LAT | | N170 AMP | |
|---|---|---|---|---|
| | LE | RE | LE | RE |
| Left eye | -11 [-16, -7] | -16 [-24, -11] | 7 [-14, 22] | -10 [-31, 17] |
| | *-0.79* | *-0.77* | *0.16* | *-0.16* |
| | *[-0.97, -0.56]* | *[-0.92, -0.52]* | *[-0.24, 0.58]* | *[-0.54, 0.20]* |
| Right eye | -11 [-18, -3] | -7 [-12, -4] | -40 [-64, -3] | 2 [-17, 14] |
| | *-0.48* | *-0.71* | *-0.41* | *0.03* |
| | *[-0.80, -0.11]* | *[-0.92, -0.44]* | *[-0.76, -0.02]* | *[-0.39, 0.41]* |

**Table S5** N170 amplitude modulation by eye visibility.

N170 amplitude modulation for left and right eye visibility, at left (LE) and right (RE) electrode. Amplitude differences are expressed as proportion of the 1st bin ERP amplitudes, i.e. amplitude modulation of 137% means that amplitude of the 10th bin ERPs was 137% the size of the amplitude of the 1st bin ERPs. Square brackets indicate 95% confidence intervals.

| | Young | | Older | |
|---|---|---|---|---|
| | LE | RE | LE | RE |
| Left eye | 137% | 153% | 129% | 161% |
| | [125, 150] | [132, 175] | [111, 150] | [141, 181] |
| Right eye | 153% | 133% | 191% | 131% |
| | [140, 166] | [124, 142] | [157, 225] | [113, 149] |

**Table S6** Effect size estimates for group differences in N170 latency and amplitude.

Effect size estimates for group differences (young – older) in N170 latency (LAT; expressed in milliseconds) and amplitude (AMP; expressed in μV/cm$^2$). Values correspond to the median of all pairwise differences between each young and every older participant. Negative latency differences indicate delays in older participants compared to young participants. Positive amplitude differences indicate larger N170 in older than young participants. A corresponding Cliff's *delta* is shown in italics. Square brackets indicate 95% confidence intervals.

| Face | Noise | Face-Noise |
|---|---|---|
| Trials without Bubbles | | |
| | N170 LAT | |
| -18 [-24, -9] | -23 [-39, -9] | 6 [-3, 16] |
| *-.64 [-.91, -.32]* | *-.51 [-.82, -.16]* | *.27 [-.13, .62]* |
| | N170 AMP | |
| .09 [-.12, .24] | .45 [.24, .64] | -.34 [-.48, -.15] |
| *.16 [-.24, .51]* | *.74 [.46, .95]* | *-.78 [-.96, -.54]* |
| Trials with Bubbles | | |
| | N170 LAT | |
| -22 [-32, -9] | -18 [-33, -6] | -2 [-8, 3] |
| *-.59 [-.86, -.26]* | *-.52 [-.82, -.20]* | *-.13 [-.50, .26]* |
| | N170 AMP | |
| -.18 [-.37, .03] | -.04 [-.18, .13] | -.15 [-.24, -.05] |
| *-.32 [-.65, .07]* | *-.11 [-.50, .26]* | *-.62[-.85 -.34]* |

### Supplementary Figures

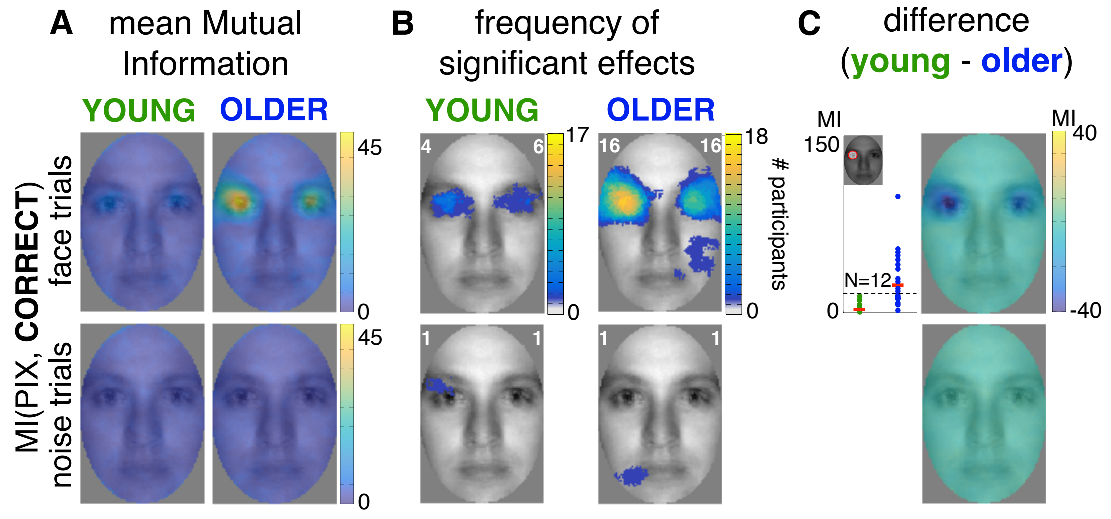


**Figure S1** Age-related differences in behavioural classification images.

**(A) Group-average MI maps** (units: scaled MI, see *Materials and Methods*). Images show average classification images for young and older participants, for face and noise trials separately. **(B) Frequency of significant effects**. The white number in the left upper corner of every image corresponds to the maximum number of participants showing a significant effect at the same pixel, whereas the number in the right upper corner corresponds to the total number of participants showing significant effects at any pixel. Eye region pixels were strongly associated with correct responses in only a few young participants and almost all older participants, suggesting that young participants used any feature to do the task, in contrast to older participants who needed to see the eye region to correctly detect a face. This reliance on the eyes was confirmed in **(C) the average classification image of the difference between the groups**. Images on the right display the differences between young and older average MI maps for face and noise trials. Scatterplots show individual MI values averaged within the left eye mask (represented as a red circle in the face inset; for explanation, see *Methods*). Red bars correspond to medians across participants. Distributions of individual MI values were different between the groups, with 12 older participants showing stronger MI than the maximum across young participants.

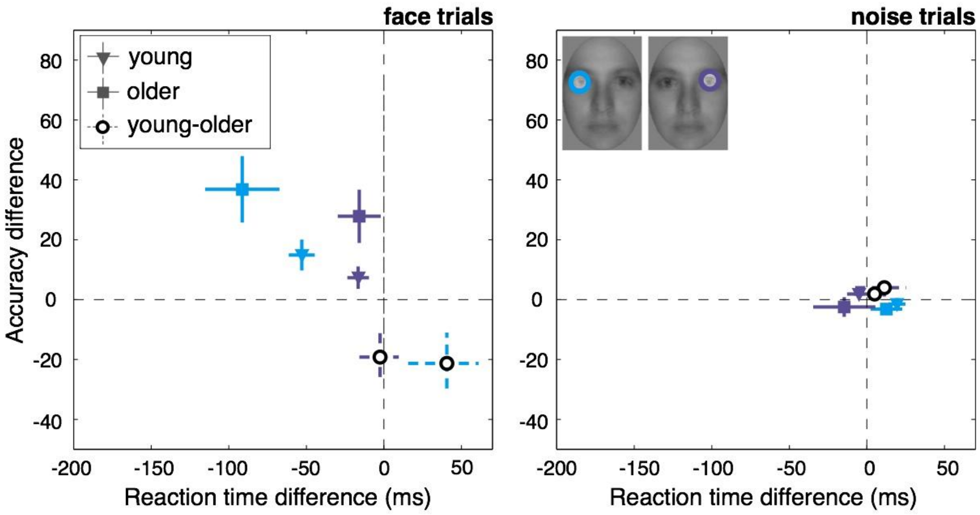


**Figure S2** Behavioural modulation by eye visibility.

Each point corresponds to the median difference between high and low visibility of the left (blue) and the right (purple) eye, separately for young (triangles) and older (squares) participants, and for face (left panel) and noise (right panel) trials. Vertical and horizontal lines mark 95% confidence intervals (CI) for accuracy and reaction time differences, respectively. Group differences (empty circles and dashed CI lines) show stronger modulation of accuracy scores by the presence of each eye in older participants, as well as stronger RT modulation by the presence of the left eye.

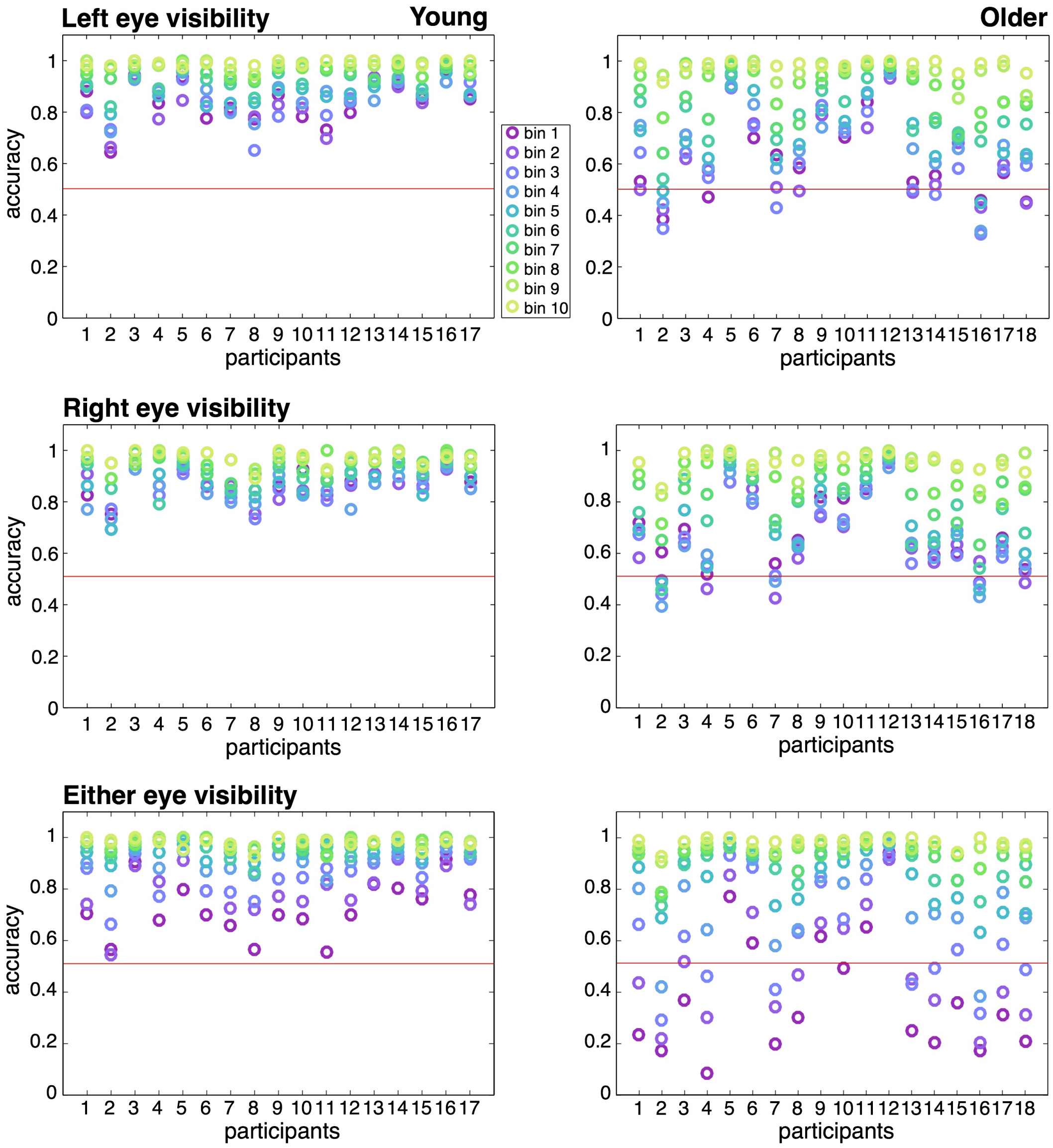


**Figure S3** Reverse analysis, face trials.

Each column of dots presents one participant's accuracy scores averaged within each of the 10 bins of visibility of the left eye (top), the right eye (middle), or either eye (bottom) from bin 1 (low visibility), to bin 10 (high visibility). Young (left) and older (right) participants' scores are presented separately. Low eye visibility (low bin numbers and purple to blue colours) was associated with lower accuracies. This association was particularly visible across older participants when there was no eye visibility in either eye region (bottom right panel). Red horizontal lines represent chance level.

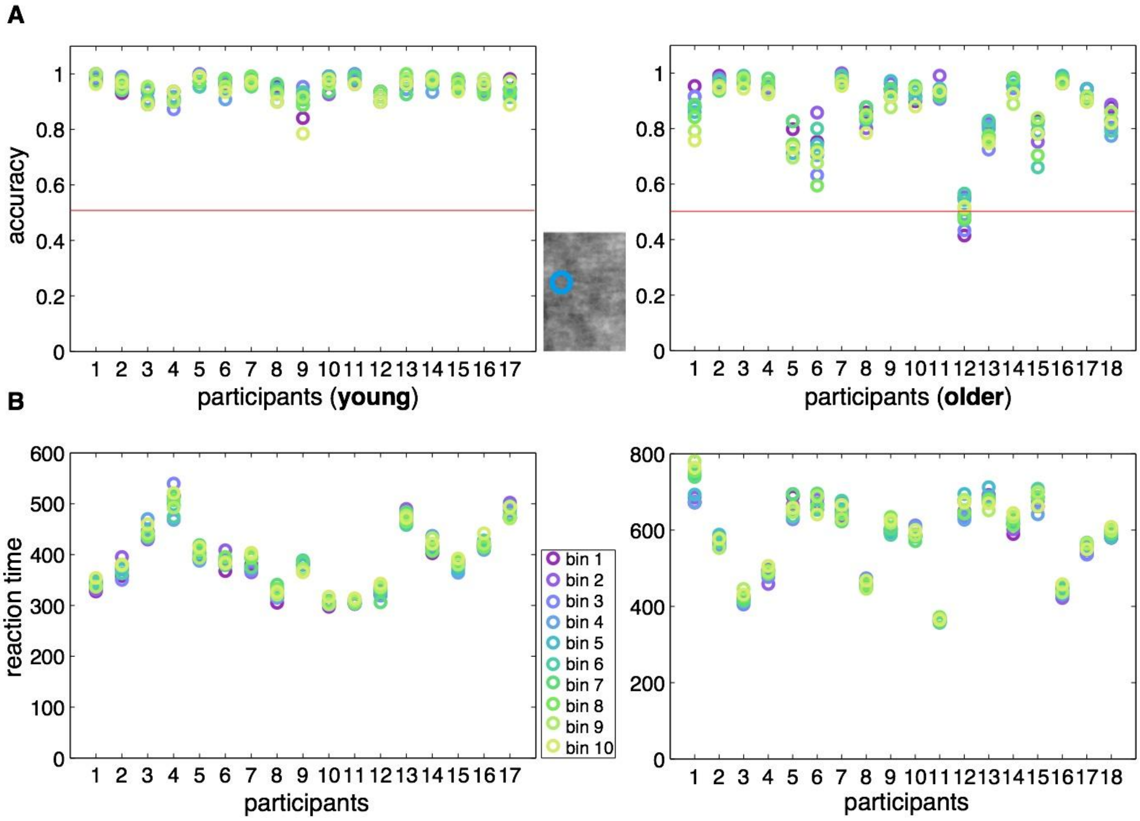


**Figure S4** Reverse analysis: left eye aperture, noise trials.

Each column of dots presents one participant's **(A)** accuracy scores and **(B)** reaction times averaged within each of the 10 bins of pixel visibility in the left eye's aperture on noise trials (bin 1: low visibility, bin 10: high visibility). Young (left) and older (right) participants' scores are presented separately. Red horizontal lines in accuracy plots represent chance level.

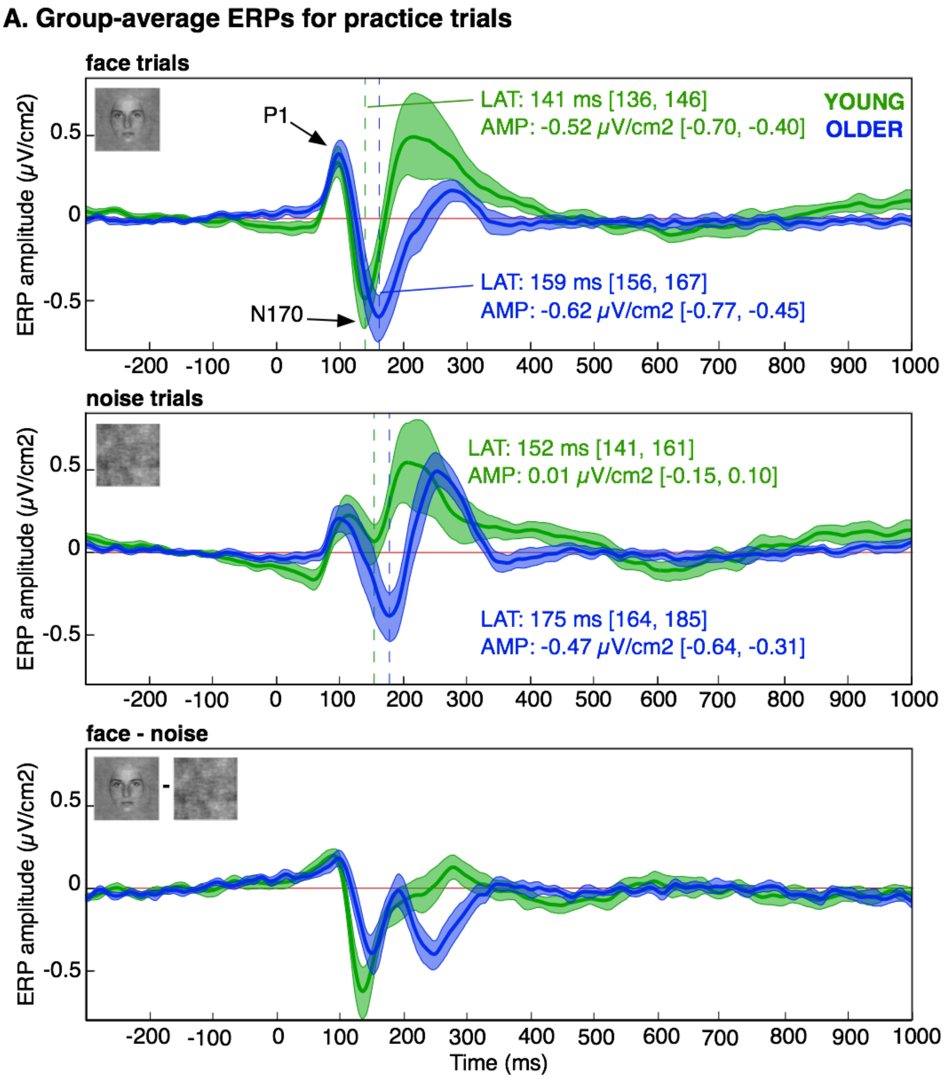


**Figure S5** Group-average ERPs for practice trials.

Thick lines correspond to ERPs averaged across young (green) and older (blue) participants, for face and noise trials separately, and for the difference between face and noise trials (third panel). Shaded areas correspond to 95% confidence intervals. Values reported in the panels correspond to median latencies and amplitudes of the N170 component.

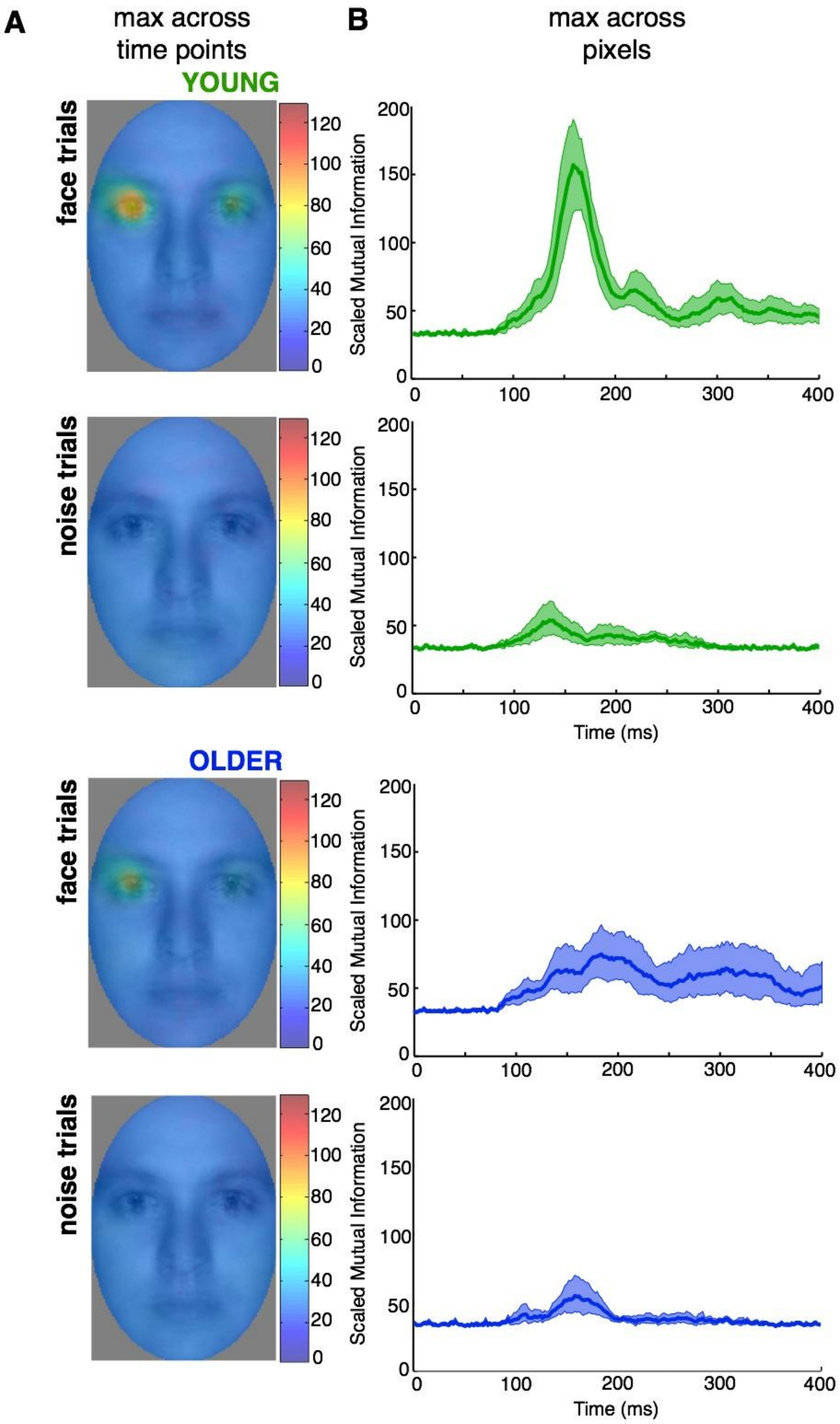


**Figure S6** MI(PIX, ERP): maximum across electrodes.

To ensure that our analysis at posterior lateral electrodes did not miss local maxima somewhere else on the scalp, we also compared between groups the maximum across electrodes of MI(PIX, ERP). The comparison was done between 0 and 400 ms following stimulus onset, by considering the maxima across pixels. We also compared the classification images by considering the maxima across time points. **(A)** Mean Classification images across participants, showing the maximum MI values across all time points and electrodes, for face and noise trials in young participants (top two images), and in older participants (bottom two images). In both young and older observers, the left eye area strongly modulated single-trial face ERPs. No clear association was seen in noise trials. **(B)** Time courses of the maximum MI across all electrodes and pixels, averaged across young and older participants, for face and noise trials separately. These time courses were very similar to those obtained in young and older participants in the main analysis, which considered only 2 lateral electrodes. Shaded areas around the time courses correspond to 95% confidence intervals.

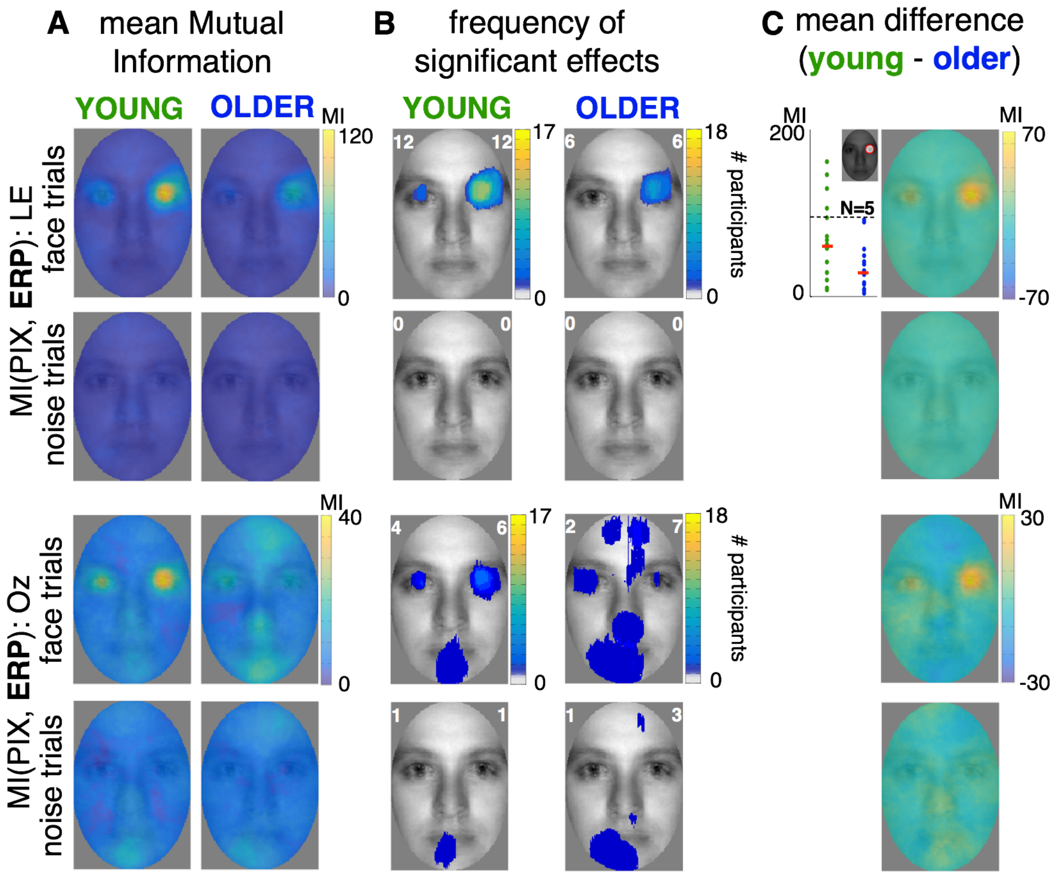


**Figure S7** Age-related differences in ERP information content.

**(A) Mean Mutual Information.** Group-average MI maps for young (left) and older (right) participants, displayed for the left (LE) lateral occipital-temporal electrode, and the midline occipital electrode (Oz) independently for face and noise trials. The classification images for face and noise trials show maximum MI values across time points. **(B) Frequency of significant effects**. The white number in the left upper corner of every image corresponds to the maximum number of participants showing a significant effect at the same pixel, whereas the number in the right upper corner corresponds to the total number of participants showing significant effects at any pixel. **(C) Differences in mean MI between young and older participants**. The scatterplot to the left of the image shows individual MI values averaged within the right eye mask. The number in each scatterplot corresponds to the number of young participants whose MI values were greater than the maximum MI value across older participants (marked as a black dashed line). The image on the right displays the difference between average young and older MI maps for every condition.

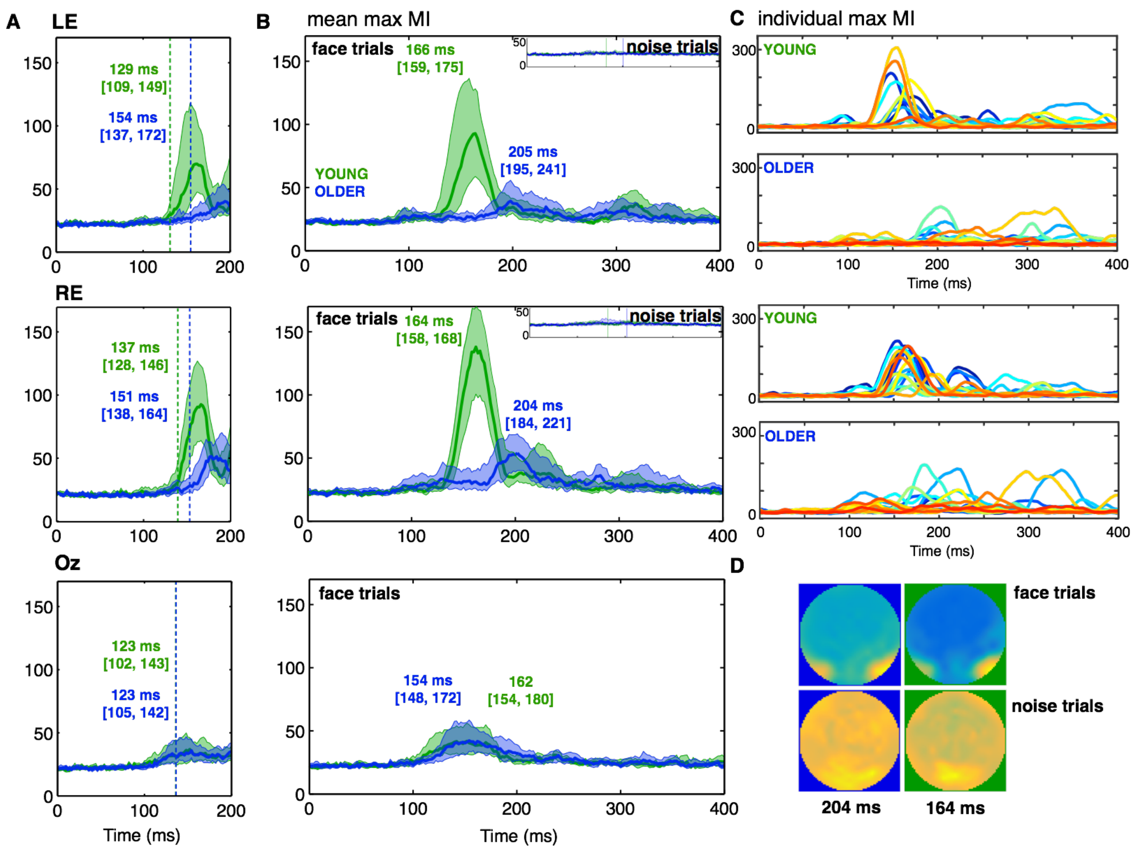


**Figure S8** Time-courses of the maximum MI across pixels.

**(A) Causal-filtered data.** Time-courses of average MI values are presented for young (green) and older (blue) participants, for face trials only. The vertical lines mark the onset of the group effect. **(B) Non-causal-filtered data.** Time-courses of average MI values are presented for both face and noise (insets) trials. Colour-coded numbers correspond to median latencies of maximum MI in both groups, obtained for face trials. **(C)** Individual participants' time-courses. In all graphs, shaded areas correspond to 95% confidence intervals around the 20% trimmed mean. **(D) Group-averaged topographic maps.** Whole-scalp MI was strongest at posterior-lateral electrodes, and tended to be right lateralised in both groups (lateralisation index for face trials, young = -0.18 [-0.31, -0.05]; older = -0.23 [-0.37, -0.09]; group difference = 0.07 [-0.07, 0.21]).

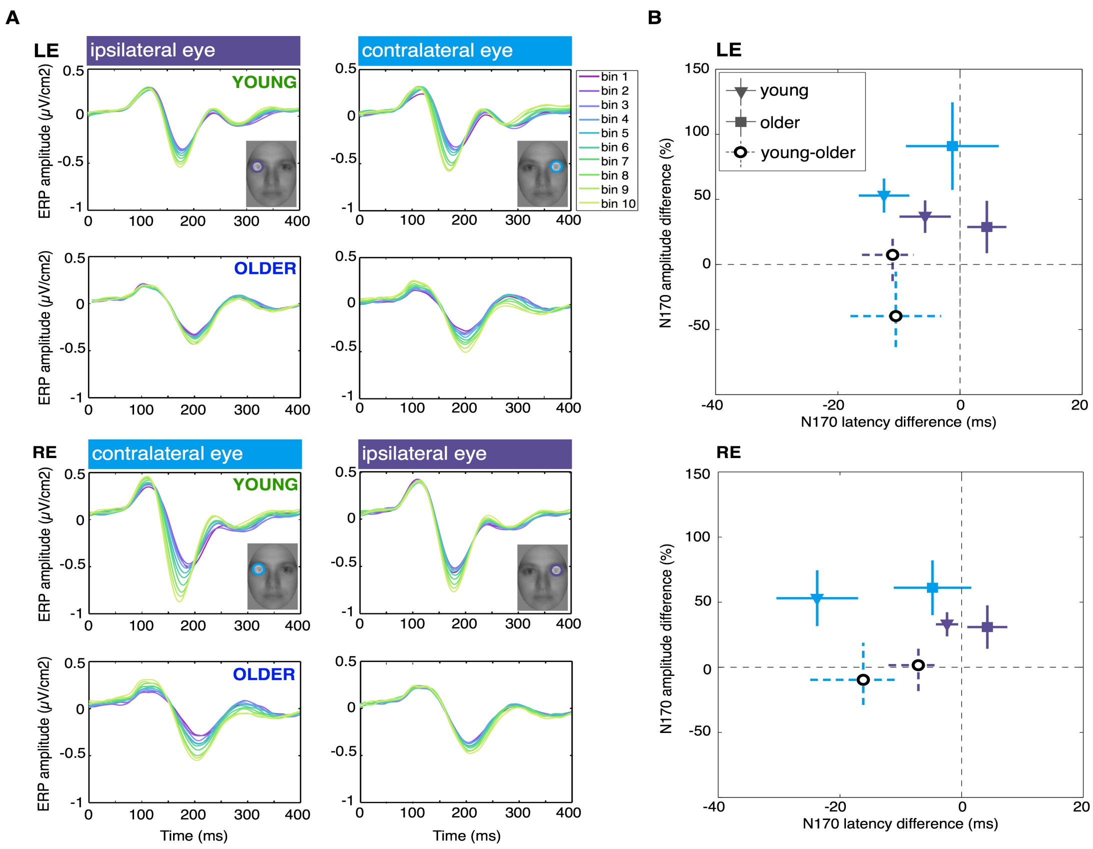


**Figure S9** ERP modulation as a function of eye visibility in face trials.

**(A) Binned ERPs**. Rows correspond to face trials in young and older participants at the left electrode (top two), and at the right electrode (bottom two). Columns correspond to ERP modulations as a function of the visibility of the contralateral eye (blue) or the ipsilateral eye (purple). In young, but not in older participants, presence of the contralateral eye was associated with earlier and larger N170, particularly at the right electrode. **(B) Quantification of eye visibility effects on the N170 latency and amplitude**. Presented are effects of eye visibility (differences between the $10^{th}$, high information, and the $1^{st}$, low information, bin) on the N170 latency and amplitude, at the left (top) and right (bottom) lateral electrodes. Amplitude and latency modulations by the presence of the contralateral eye (blue) and ipsilateral eye (purple) are presented in both plots. Amplitude differences are expressed as proportion of the $1^{st}$ bin ERP amplitudes: an amplitude difference of 50% means that amplitude of bin 10 ERPs was 150% the size of the amplitude of bin 1 ERPs. Filled circles correspond to median ERP modulations across young participants; squares show medians across older participants; empty circles show differences between group medians. Vertical and horizontal bars correspond to 95% confidence intervals.

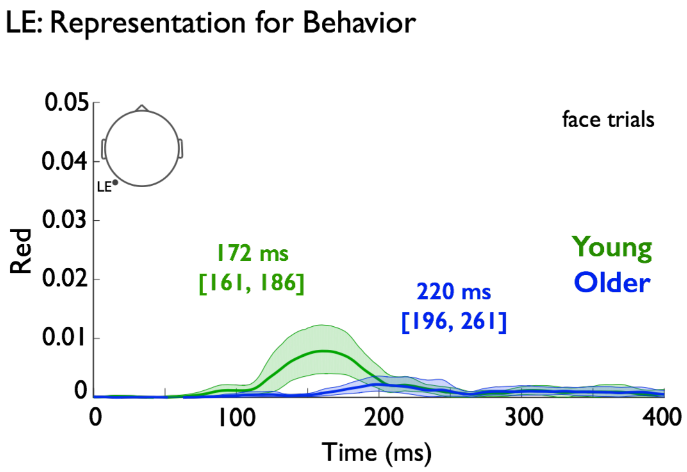


**Figure S10** LE: Representation for behaviour.

Time course of behavioural redundancy showing a 42 ms representation of the right eye delay on RE for face detection reaction time in older adults.

### Supplementary References


1. Smith ML, Gosselin F, Schyns PG. Receptive fields for flexible face categorizations. Psychol Sci. 2004;15(11):753–61.

2. Rousselet GA, Ince RAA, van Rijsbergen NJ, Schyns PG. Eye coding mechanisms in early human face event-related potentials. 2014;14:1–24.

3. Cliff N. Ordinal methods for behavioural data analysis. Mahwah, NJ: Lawrence Erlbaum Associates; 1996.

4. Wilcox RR. Graphical Methods for Assessing Effect Size: Some Alternatives to Cohen's d. J Exp Educ. 2006;74(4):353–67.

5. Wilcox RR. Introduction to robust estimation and hypothesis testing. Academic Press; 2012. 690 p.

6. Ince RAA, Giordano BL, Kayser C, Rousselet GA, Gross J, Schyns PG. A statistical framework for neuroimaging data analysis based on mutual information estimated via a gaussian copula. Hum Brain Mapp. 2017 Mar;38(3):1541–73.

7. Nelsen RB. An Introduction to Copulas. New York, NY: Springer New York; 2006. (Springer Series in Statistics).

8. Cremer R, Zeef EJ. What Kind of Noise Increases With Age? J Gerontol. 1987;42(5):515–8.

9. Salthouse TA, Prill KA. Effects of aging on perceptual closure. Am J Psychol. 1988;101(2):217–38.

10. Whitfield KE, Elias JW. Age cohort differences in the ability to perform closure on degraded figures. Exp Aging Res. 1992;18(2):67–73.

11. Kurylo DD. Effects of aging on perceptual organization: efficacy of stimulus features. Exp Aging Res. 2006;32(2):137–52.

12. Roudaia E, Bennett PJ, Sekuler AB. The effect of aging on contour integration. Vision Res. 2008;48(28):2767–74.

13. Danziger WL, Salthouse TA. Age and the perception of incomplete figures. Exp Aging Res. 1978;4(1):67–80.

14. Lindfield KC, Wingfield A. An experimental and computational analysis of age differences in the recognition of fragmented pictures: inhibitory connections versus speed of processing. Exp Aging Res. 1999;25(3):223–42.

15. Lindfield KC, Wingfield A, Bowles NL. Identification of fragmented pictures under ascending versus fixed presentation in young and elderly adults: Evidence for the inhibition-deficit hypothesis. Aging, Neuropsychol Cogn. 1994;1(4):282–91.

16. Salthouse TA, Lichty W. Tests of the neural noise hypothesis of age-related cognitive change. J Gerontol. 1985;40(4):443–50.
